# Evaluating the accuracy of the umbrella sampling plots with different dissociation paths, conformational changes, and structure preparation

**DOI:** 10.1101/169532

**Authors:** Wanli You, Zhiye Tang, Chia-en A. Chang

## Abstract

The kinetics of ligand dissociation has been found to be crucial for a good drug candidate. Therefore, examining the underlying free energy profile of the dissociation that governs the kinetics becomes important. Umbrella sampling (US), a widely used free energy calculation method, has long been used to explore the dissociation process of ligand-receptor systems. The potential of mean force (PMF) computed from US seems to always produce binding affinity and energy barriers that more or less agree with experiments. However, such PMFs are influenced by many practical aspects, like the method used to generate the initial dissociation pathway, collective variables (CVs) that used to describe the reaction coordinate (RC), and how intensive the sampling is in the conformational space restrained by the CVs. These critical factors were rarely studied. Here we applied US to study the dissociation processes of β-cyclodextrin (β-CD) and p38α complex systems. For β-CD, we used three different β-CD conformations to generate the dissociation path manually. For p38α, we generated the dissociation pathway using accelerated molecular dynamics (AMD) followed by conformational relaxing with short conventional molecular dynamics (MD), steered molecular dynamics (SMD) and manual pulling. We found that even for small β-CD complexes, different β-CD conformations will alter the height of the PMF and different dissociation directions result in appearance/disappearance of local minima. SMD poorly samples the residue sidechain movement, leading to overestimated height of PMF. On the other hand, the AMD pathway relaxed by short conventional MD sampled more accurate structures, resulting in reasonable PMF.

## Introduction

Free energy is an important quantity that characterizes chemical and biological processes. The change of free energy governs the directionality and extent of chemical reactions. Free energy decomposes into enthalpy and entropy, where determination of entropy is challenging both experimentally and computationally. For these reasons, it is one central task for computational chemist to achieve accurate calculation of free energy, especially the free energy profile along a chemical process [1–3]. A variety of free energy calculation methods have been developed in the past decades, such as perturbation theory [4], thermodynamic integration [5], umbrella sampling [6], and partition function from density of states [7], and provided insights into various chemical and biological systems [8, 9]. Among these methods, umbrella sampling (US) is a conceptually straightforward, computationally efficient and reliable one that computes the potential of mean force (PMF) based on rigorous probability calculations [6]. It requires a well-defined reaction coordinate (RC) represented by one or a few collective variables (CVs) [10]. Intensive conformational sampling is performed by enforcing external restraints at the conformations along the RC within a series of successive overlapping windows. Finally, the PMF can be constructed by removing the external restraints.

US is particularly suitable for computing the PMF of ligand-receptor dissociation driven by non-bonding interactions. It has long been applied to calculate binding affinity of various receptor-ligand systems, ranging from small chemical molecular system [11] to large biological systems [12]. Furthermore, by sampling local energy barriers along dissociation path, it can provide thermodynamic details for molecular recognition. However, US itself does not provide the dissociation path. Therefore, enhanced sampling methods, such as steered molecular dynamics simulation (SMD) [13–16], adaptive biasing force (ABF) [17] and metadanamics [18], are often used to provide the dissociation pathway that can be used as initial conformations for the intensive biased sampling for US. A number of methods have been proposed to improve the accuracy of US, such as the use the constrained schemes to alleviate sampling limitation [19–21], or the combination with other method like Markov model to improve the convergence [22]. Despite the natural connection between these methods and US, how accurately the conformations from these enhanced sampling methods can resemble true dissociation path, or whether they can provide reasonable initial conformations for US, have rarely been studied. Moreover, how one dissociation path compared to another in the dissociation pathway ensemble may affect the results from US remains unclear. Here we answered these two questions by studying influential factors of US using two systems, β-Cyclodextrin (β-CD)-ligand complexes and p38α-inhibitor system.

β-CD is a cyclic oligosaccharide containing seven glucopyranose units linked through 1,4 α glycosidic bonds, thus forming a truncated conical structure. With a hydrophobic inner surface and hydrophilic rims containing primary and secondary hydroxyl groups, it is able to accommodate small hydrophobic molecules, therefore enhancing the solubility and bioavailability of such molecules (Figure 1). The cavity of β-CD also resembles a protein binding site, and this makes it a good host molecule to study ligand-receptor binding. Because of these properties, β-CD and its derivatives have been widely used in drug delivery, pharmaceutical, food, and chemical industries [23–29]. Therefore, a large amount of experimental reference data is available [30–37].

**Figure 1.**
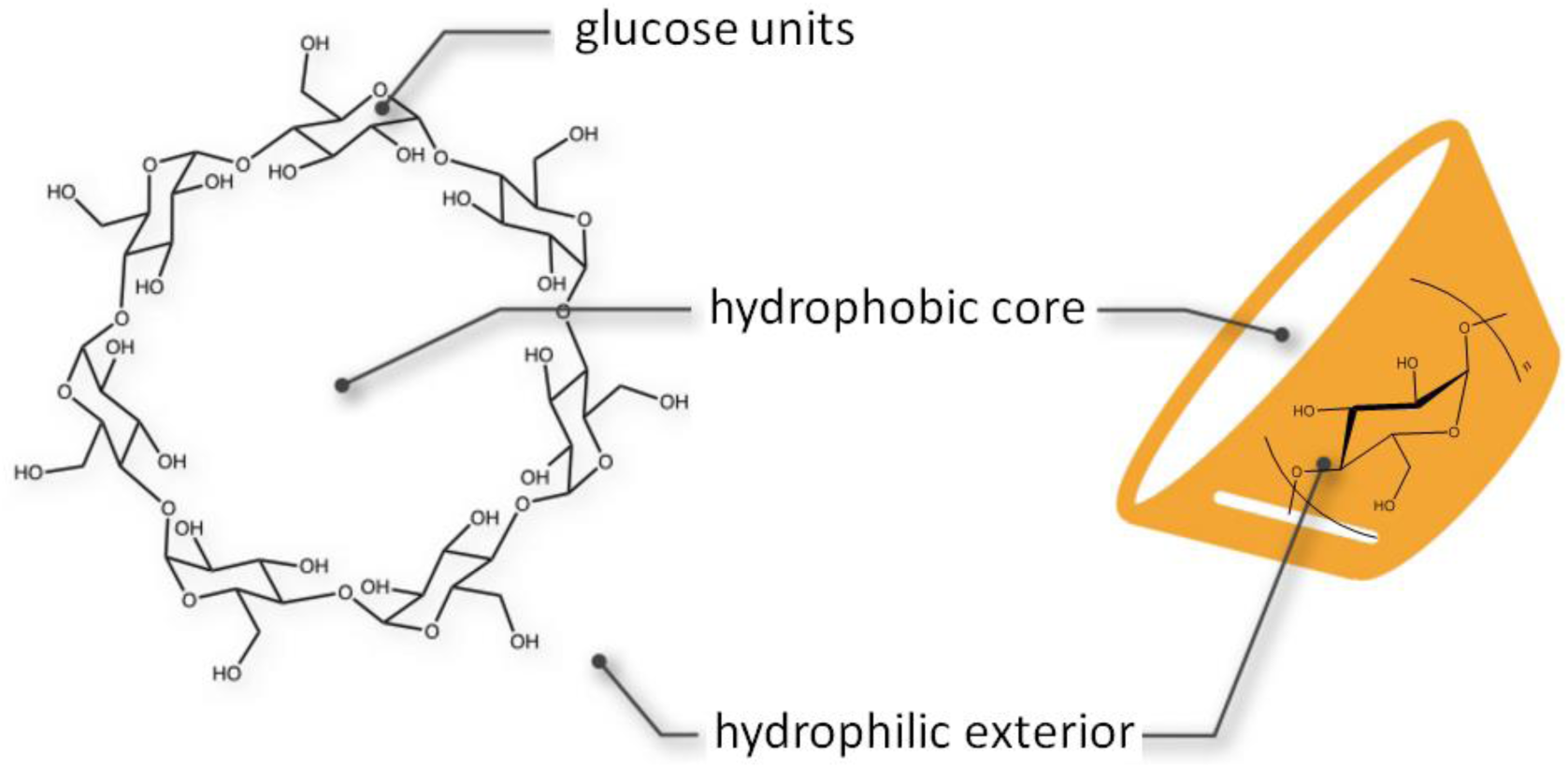
Structure of β-cyclodextrin (β-CD). β-CD is consisted of seven glucopyranose units with a hydrophobic core and hydrophilic exterior.

p38α is the major isoform of p38, which belongs to the mitogen-activated protein kinases (MAPKs), a super-family of enzymes that regulate a variety of biological processes, such as proliferation, gene expression, differentiation and apoptosis [38, 39]. p38α has been a drug target for treating various inflammatory diseases, including rheumatoid arthritis, asthma, and cardiovascular disease [40–42]. To develop new inhibitors, it is necessary to investigate the kinetics behaviors of inhibitors during dissociation process. Like all protein kinases, p38α consists of a N-terminal domain and a C-terminal domain that are connected via a hinge [43]. The activation loop that carries a DFG (Asp-Phe-Gly) motif determines the opening or the closing of the binding cavity, where ATP binds during the activation process. The conformational change of activation loop can be characterized by different orientations of its Phe sidechain. In the active conformation, Phe is buried in αC helix (DFG-in), while in inactive conformation, Phe rotates away from the αC helix and projects into the ATP binding pocket (DFG-out). SB2 is a ligand of p38α that can bind to ATP binding site while activation loop adopts either DFG-in or DFG-out conformation [44] (Figure 2) and doesn’t interfere with the conversion between DFG-in and DFG-out conformations of activation loop. This makes it the perfect candidate to study the influence of receptor conformational change on construction of free energy profile.

**Figure 2.**
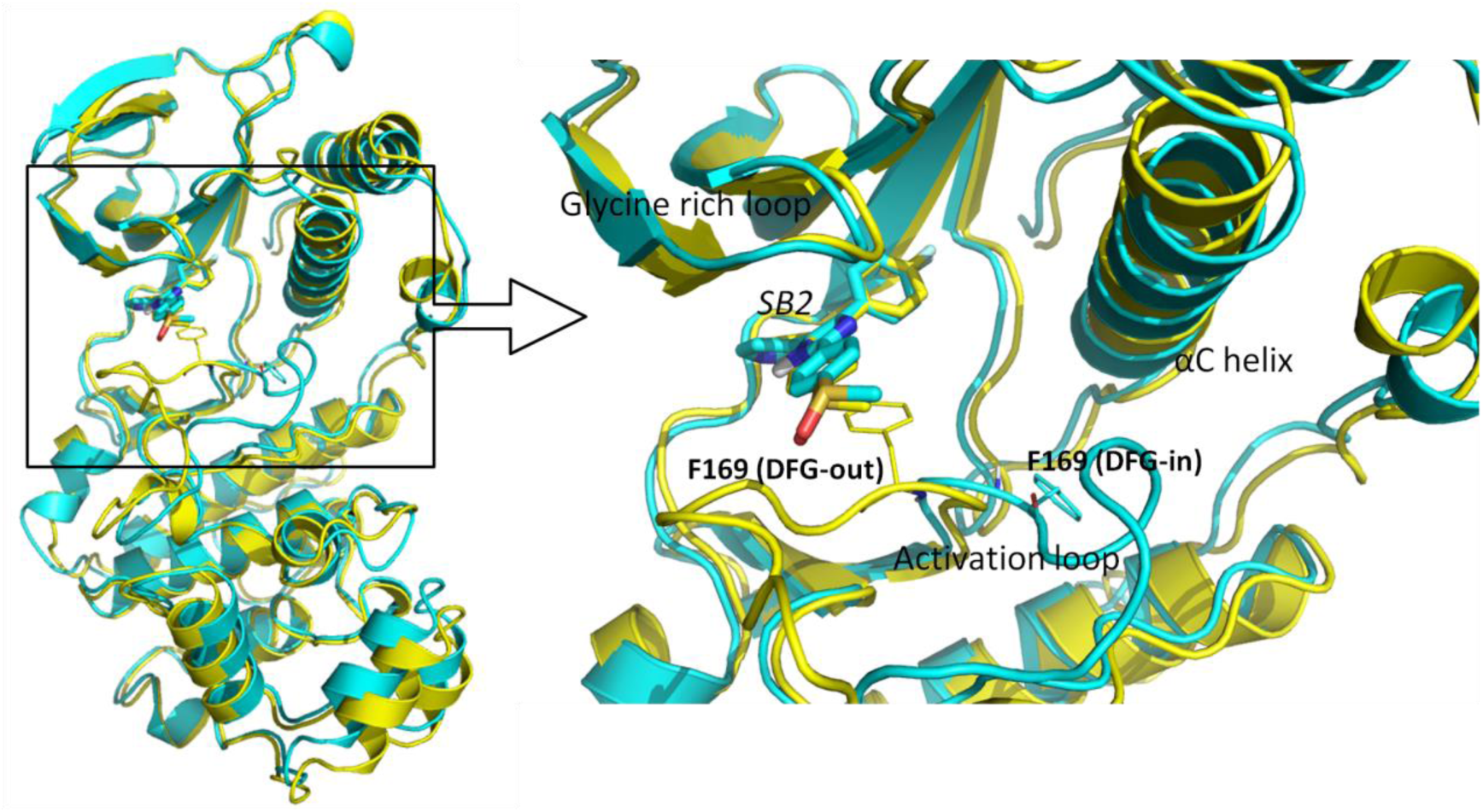
The comparison of the bound structures of SB2 in DFG-in and DFG-out conformations. The left figure shows the structure alignment of DFG-in (cyan, PDB 1A9U) and DFG-out (yellow, PDB 3GCP) conformations bound with ligand SB2. The right figure shows binding site structure of p38α-SB2 complex. The Phe169 from DFG motif is shown in thin licorice structure, ligand SB2 is shown in bold licorice structure.

In this work, we investigated how the PMF from US is affected by subtle changes in the dissociation pathways and conformational sampling methods that provide the initial conformations. We used dissociation of β-CD-aspirin, β-CD-1-butanol, and p38α-SB2 systems as examples. We utilized accelerated MD (AMD), SMD and manual pulling methods to provide the dissociation pathways as initial conformations for US, and investigated how these methods affect the PMF from US. By using different β-CD conformations as starting point for performing US, we found that the host conformation may fundamentally change the depth of the PMF from US, and the influence of initial conformation can hardly be removed by nanosecond level biased MD simulation, even for small systems like a β-CD complex. We also showed that AMD is a good tool to provide initial conformations for US simply by relaxing the conformations along dissociation pathways sampled by AMD using short MD simulations. However, SMD may not be a suitable method to provide the initial conformations, because the dissociation pathway sampled by SMD lacks important residue sidechain movement.

## Materials and methods

### Structure preparation and parameters

*β*-*cyclodextrin*. We selected three β-CD conformations from previous MD simulations of β-CD-aspirin complexes and removed the ligand as initial conformation of β-CD. Two ligands, aspirin and 1-butanol, were selected to perform US along manually built dissociation pathways as detailed in later section (Figure 3). We used q4MD-CD force field for β-CD [45]. We manually built the ligand structures with Vega ZZ [46] and computed the partial charges for them by using B3LYP/6-31+G(d,p) ChelpG calculations with Gaussian package [47] after optimizing the structures using the same settings. GAFF was used for the ligands.

**Figure 3.**
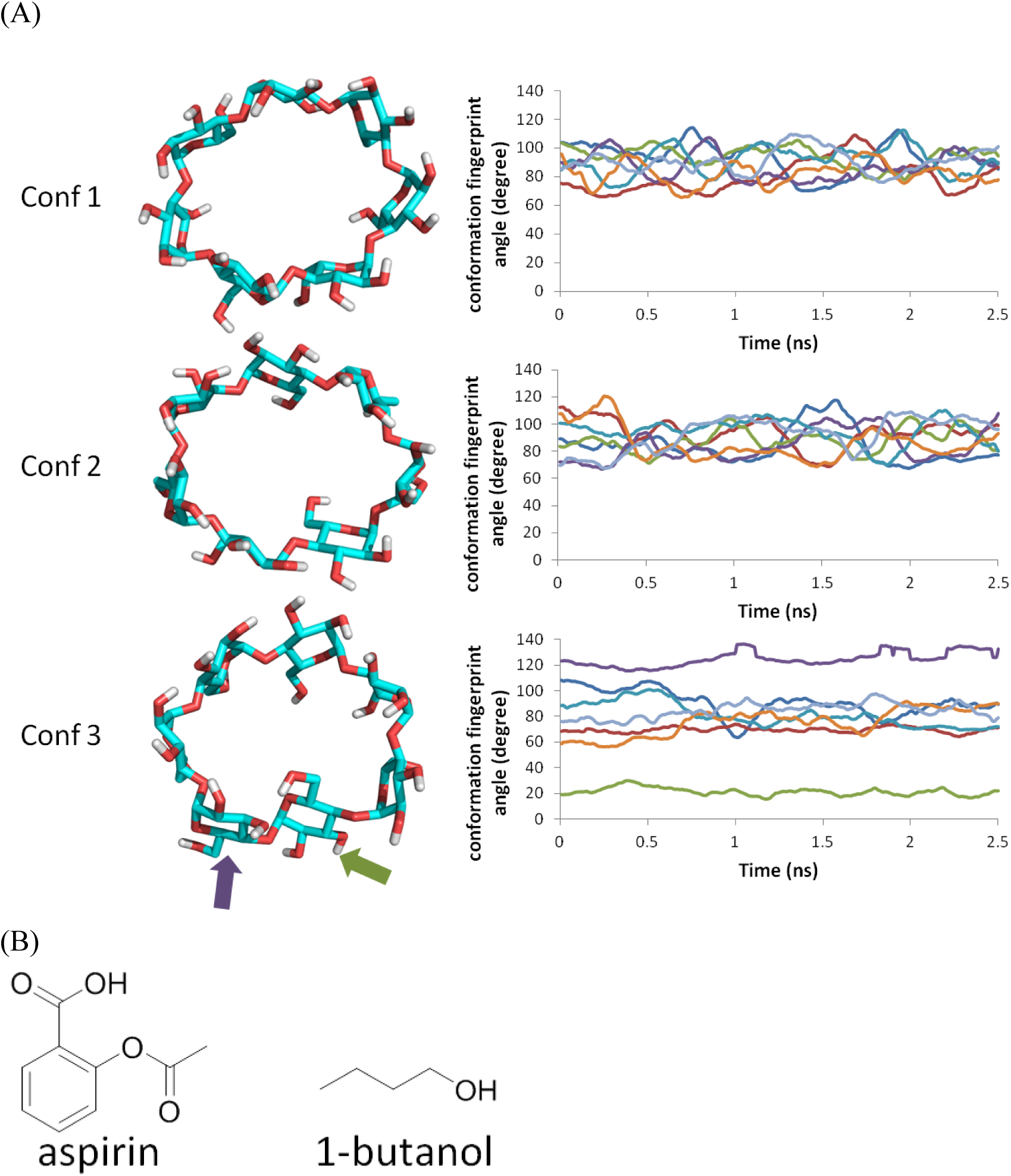
Structures of β-CD and ligands and fingerprint dihedral plot of three β-CD conformations. (A) Three different conformations of β-CD. The plots of their conformation fingerprint angles (defined in Figure S1) along trajectories are on the right. The fingerprint dihedral plots of β-CD are from biased MD of one US window where distance between center of mass (COM) of β-CD and aspirin is restrained to 10 Å. Before measurement of fingerprint angles, the trajectories were smoothed by averaging 100 forward and 100 backward frames on the concurrent frame throughout the whole trajectory to remove the noise. (B) Structures of aspirin and 1-butanol.

*p38α*. SB2 is a ligand of p38α system that binds to both DFG-in and DFG-out conformations. We obtained p38α DFG-in conformation (PDB ID: 1A9U) [48] and DFG-out conformation (PDB ID: 3GCP) [49] from protein data bank (PDB). We built the missing activation loop (residues 173 to 184) of 3GCP by using the conformation from previous MD simulation of free DFG-out p38α. Amber 99SB force field was used for proteins and GAFF was used for ligand SB2.

### Preparation of dissociation paths for US

For β-CD, we manually docked the ligands along the dissociation path. First, we put the center of mass (COM) of β-CD at the origin (0, 0, 0), and aligned its principal axes along X, Y, Z axes so that the primary cavity of β-CD faces the positive direction of X-axis. Then, we manually located the ligand so that its COM is also aligned at origin. By using this artificial bound conformation, we gradually moved the ligand along positive and negative of X-axis at a speed of 0.1 Å every step for 26 Å in both directions, until the ligand is fully dissociated. In this way, we obtained path A and B for ligand dissociation along the primary and secondary cavity of β-CD (Figure 4). We repeated this procedure for aspirin and 1-butanol in the three β-CD conformations respectively.

**Figure 4.**
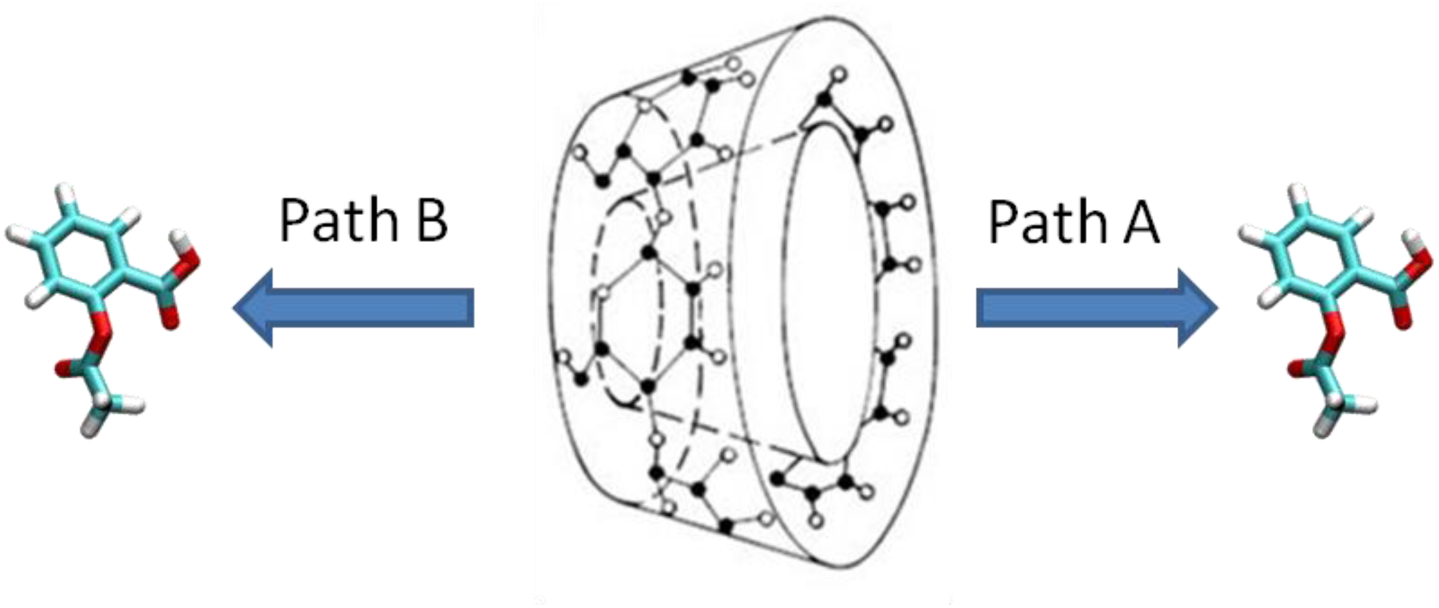
Dissociation pathways of β-CD complexes. Path A is from the primary cavity and path B is from secondary cavity of β-CD.

For p38α-SB2 system, we constructed the dissociation paths by using three ways, i.e. AMD, SMD, and manual pulling as used for β-CD. We obtained two paths from AMD (path1 and path2), one path from SMD, one path from manual pulling for DFG-in conformation, and one path from AMD for DFG-out conformation.

#### Accelerated MD simulation

AMD, which introduces a continuous non-negative bias boost potential function ΔV(r)to the potential energy surface when the system potential is below a reference energy, to enhance the conformational sampling of biological systems, therefore lowering the local barriers to accelerate the calculation [51]. AMD uses following equations to alleviate the energy barriers,

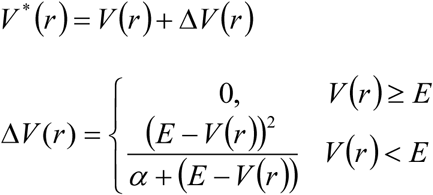

where V(r) is the original potential, E is the reference energy, and V^*^(r) is the modified potential. ΔV(r) is the boost potential, α is the acceleration factor.

The boost potential ΔV(r) can be applied to dihedral with input parameters (Ed, αD) and overall potential energy terms with input parameters (Ep, αP),

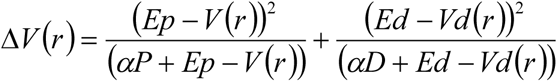

For simulations of p38α, we applied both potential-boost and dihedral-boost.

#### Steered MD simulation

SMD simulation uses a time-dependent external force to drive the system to move in a predefined way [52]. The external force V(t) can be described as,
V(t) = *k*[x-x_0_(t)]^2^
where x and x_0_(t) are the CV in simulation and the predefined time-dependent track of the CV, k is a harmonic force constant. We selected the distances between Cα of Arg73 and CC2 of SB2 (Figure S2), which is also used to describe the RC in US, as the CV. x_0_(t) was set to move by 1.75 Å/ns and with a maximum at 33 Å. The force constant k was set to 10 kcal/mol⋅Å^2^. We equilibrated the bound state conformation for 100 ns using conventional MD, and then performed SMD for 10 ns at 300K and 1 bar in NPT ensemble. Temperature was maintained by Langevin thermostat.

#### Manual pulling

In manual pulling, crystal structure of SB2 bound to p38α (1A9U) was used as reference conformation. Ligand SB2 was gradually moved along the vector of Cα of Arg73 and CC2 of SB2 towards outside of cavity at a speed of 0.25 Å every step for 16.75 Å, until SB2 was fully dissociated.

### Umbrella sampling

US [53, 54] was performed to compute the free energy along the dissociation pathway. In US, a series of windows are evenly located along RC and intensive sampling in these windows is achieved by enforcing an external biasing potential. The samplings in each window must overlap with adjacent windows, so that the unbiased PMF can be reproduced by removing the biasing potential. The external biasing potential *u_i_* at window *i* is a harmonic function *u_i_* = *k_i_(r* – *r_i_)*^2^, where *r_i_* is the reference position, and *k_i_* is the harmonic force constant. All biased MD simulations were performed with Amber14 [55]. We used WHAM [56] to remove the biasing potential and reconstruct the PMF.

For β-CD, the distance between COMs of heavy atoms of β-CD and ligand was selected as the CV to represent the RC. As described in previous section, a total of 260 windows with 0.1 Å spacing along the RC were used to performed biased MD simulation. We minimized the initial conformation for each window for 1500 steps using generalized Born (GB) implicit solvation model [57] to remove clashes from manual pulling. Then we solvated β-CD-ligand complex with a 30-Å rectangular box of TIP3P water molecules using tleap module in AMBER14 [58]. After minimizing the system for 1000 steps, we equilibrated the water molecules at 298K for 1 ns with 1-fs timestep, and heated the entire system at 200K, 250K and 298K for 150 ps. In the minimization and equilibration steps, a harmonic force constant of 1000 kcal/mol⋅ Å^2^ was used to restrain the ligand at the correct window. Finally, we performed 2.5 ns production run at 298K with restraint using harmonic force constant of 100 kcal/mol⋅Å^2^. In WHAM, the bin size was set to 0.05 Å. The tolerance for iteration was set to 0.0001. The temperature was set to 298 K.

For p38α, the distance between Cα of Arg73 and CC2 of SB2 was selected as the RC. We set up five sets of biased MD simulations for computing PMF using US as detailed in the previous section. Note that we performed 10 ns conventional MD simulations on the conformations along the two DFG-in and one DFG-out paths yielded from AMD to equilibrate the conformations before we used them as initial conformations in each window for the biased simulations for US (Figure S2). From the SMD trajectory and the conventional MD that used to relax conformations of AMD, we selected the conformations that fall into the windows along RC and have minimal SB2 root mean square deviations (RMSD) compared to the bound state SB2 as initial conformations for the corresponding window. For the manual pulling path, since the system is not optimized or solvated, we performed these following steps on the conformations we obtained before proceeding to biased MD sampling. We optimized the conformations along the path by minimizing the hydrogen atom, sidechains, and entire complex for 500, 5000 and 5000 steps respectively for the same reason as we did for β-CD. Next, we solvated the conformation using TIP3P water model [58] so that the edge of the water box is at least 12 Å away from the solutes. We also added Na^+^ ions to neutralize the system. We optimized the water molecules and the entire system for 10000 and 20000 steps respectively. After equilibrating the solvate for 40 ps at 298K in NPT, we heated the system from 250K to 300K gradually. In the minimization and equilibration process, the external force constant was 500 kcal/mol⋅Å^2^. In total, 62 windows from SMD, 68 windows from path 1, 2, manual pulling, and 71 windows from DFG-out path were evenly located every 0.25 Å along the RC. For all five sets of conformations, we performed a production run for 10 ns at 300K with an external restraint of 5 kcal/mol⋅Å^2^. In WHAM, the bin size was set to 0.2 Å. The tolerance for iteration was set to 0.0001. The temperature was set to 300 K.

## Results and Discussion

### Unbinding process of β-CD complex system

The PMFs of path A and B are constructed by using WHAM from bound state to free state, and combined so that the free states of two paths of have the same free energy. The combined PMFs of β-CD-aspirin and β-CD-1-butanol are shown in Figure 5. Comparing Conf 1 and 2 with Conf 3 of the β-CD complex systems, it’s clear that the host conformations have remarkable impact on the shape of the PMF. For aspirin, the binding affinities of Conf 1 and 2 are −2.8 and −2.7 kcal/mol which are similar, while the binding affinity of Conf 3 is only −1.8 kcal/mol. The binding affinities from Conf 1 and 2 are 1 kcal/mol less favorable than experimental value −3.77 kcal/mol. For 1-butanol, the binding affinities of Conf 1, 2 and 3 are −1.7, −1.7, −1.3 kcal/mol, respectively. The binding affinities of Conf 1 and 2 agree with experimental value (−1.67 kcal/mol). We only considered three β-CD conformations but in reality, β-CD can adopt much more conformations in the ligand association and dissociation. Also, in the biased MD simulations, the external harmonic potential was only applied on the direction of the RC which was represented by the COM distance CV, and the ligand is free to move on the sphere with a radius of the COM distance in that window, resulting in unrestrained deviation from the X-axis. For these two reasons, the computed binding affinities do not rigorously agree with the experimental values.

**Figure 5.**
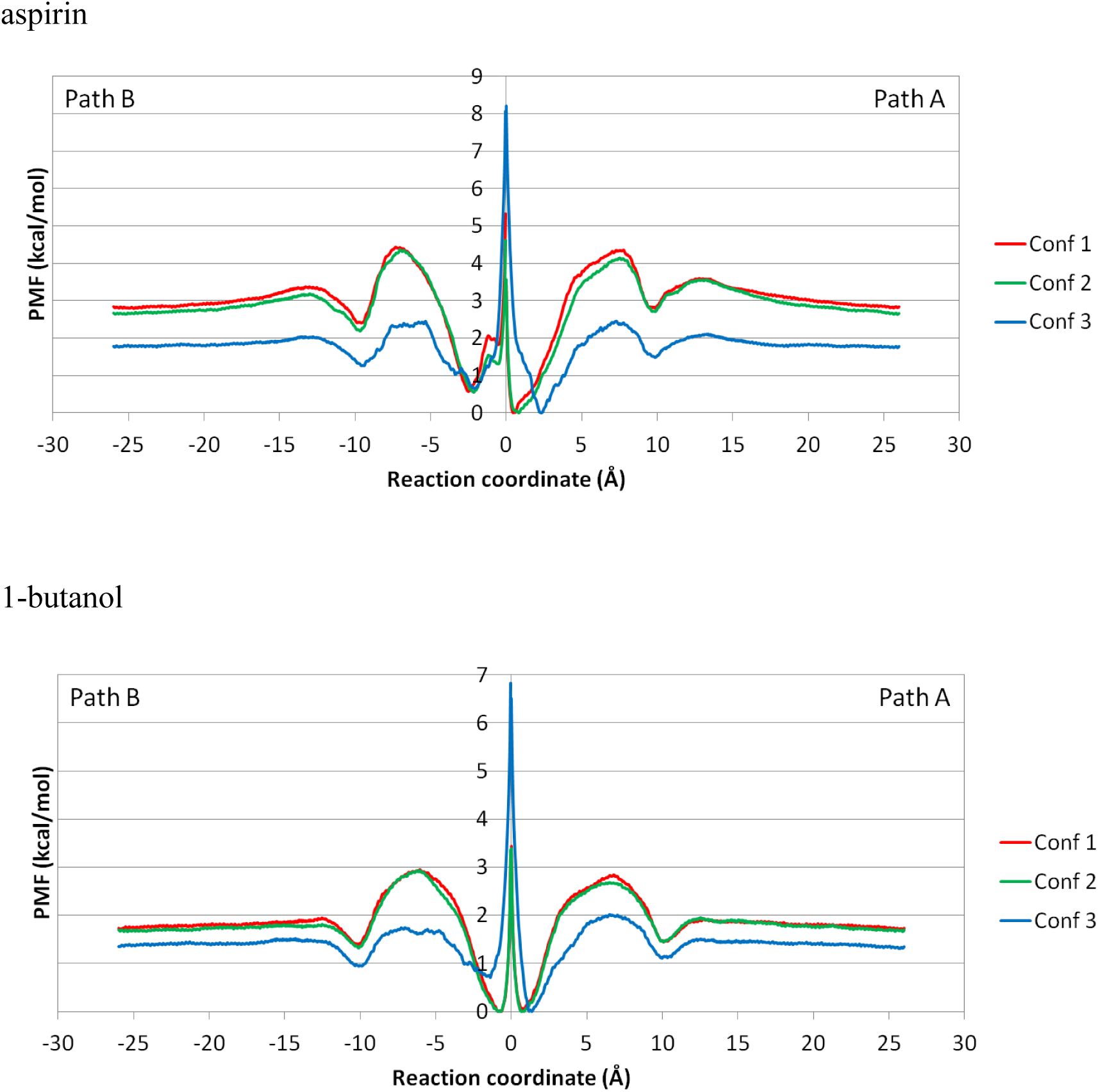
Combined PMF of dissociation of aspirin and 1-butanol from β-CD. Path A and Path B are combined for Conf 1, 2 and 3 of β-CD-aspirin and β-CD−1-butanol complexes.

The PMFs (Figure 5) also suggest that the association energy barriers of aspirin and 1-butanol are 1.5 and 1.1 kcal/mol in Conf 1 respectively. The similar association energy barriers of aspirin and 1-butanol agree with the fact that these two ligands have similar association rate constants (Table 1). Although the two-fold faster association rate constant of aspirin implies a smaller association energy barrier and the computed value is actually bigger than 1-butanol, the difference in the barrier is small than 0.6 kcal/mol and can be considered as bias from the β-CD conformation and errors due to thermal fluctuation. The computed dissociation energy barriers of aspirin and 1-butanol in Conf 1 are 4.4 and 2.8 kcal/mol respectively. This agrees with the experimental dissociation rate constants perfectly.

**Table 1.** Experimental kinetics and thermodynamics data of aspirin, 1-butanol complexed with β-CD, and SB2 complexed with p38α. *K*_D_, *k*_on_ and *k*_off_ data for complexes were taken from [31, 34, 50], Δ*G*_exp_ of β-CD−1-butanol complex was taken from [35], Δ*G*_exp_ of β-CD-aspirin and p38α-SB2 complexes were calculated using Δ*G*_exp_ = RT ln *K*_D_.

| Complex system | $K_D$<br>(nM) | $k_{on}$<br>[M<sup>-1</sup>s<sup>-1</sup>] | $k_{off}$<br>[s<sup>-1</sup>] | $\Delta G_{exp}$<br>(kcal/mol) |
| --- | --- | --- | --- | --- |
| β-CD-aspirin | $1.81 \times 10^6$ | $7.20 \times 10^8$ | $1.30 \times 10^6$ | -3.77 |
| β-CD-1-butanol | $1.36 \times 10^8$ | $2.80 \times 10^8$ | $3.80 \times 10^7$ | -1.67 |
| p38α-SB2 | 11.5 | $1.5 \times 10^7$ | $1.8 \times 10^{-1}$ | -10.90 |

The combined PMF unambiguously indicates that the ligands bind preferentially to the primary direction of the β-CD cavity. The ligand binds to β-CD with a range from −3 to +3 Å in depth, and there is a huge energy barrier in all combined PMFs within this range. By close investigation of the population plot (Figure S3), we noticed that this energy barrier near origin of the RC was caused by abnormal behavior of COM distance restraint when the two COMs were close to each other (**SI Section Artifact of Restraints**). Therefore, this energy barrier at the origin of the RC can be ignored. By looking at the combined PMFs without this energy barrier, we still observe that the free energy of the primary side is consistently 0.6 kcal/mol more favorable than the secondary side with one exception of 1-butanol in Conf 1 and 2. This is because the primary side of the β-CD cavity is more open and allows the aspirin to better fit into it. Because of its relatively small size, 1-butanol can fit into both primary and secondary sides of the β-CD cavity in Conf 1 and 2. In Conf 3, the size of cavity shrinks due to the flipping of two of the glucopyranose units of β-CD, 1-butanol prefers to bind to the bigger primary side of β-CD cavity (Figure 3).

Since we only put restraints on the 1-D RC represented by the COM distance CV, we don’t have control on the position of the ligand on the sphere centered at the COM of β-CD. For example, in the window where the COM distance is 10 Å, the ligand can adopt positions anywhere on a sphere with a radius of roughly 10 Å if the interactions between the ligand and β-CD is not considered. This is not a problem in the ideal case where interactions are ignored, but in reality, the intermolecular attractions may alter the distribution of ligand on such a sphere and significantly deviates the ligand from the artificial dissociation path along the X-axis (Figure 6). In our simulation for aspirin, the ligand more or less follows the X-axis dissociation path way within 7 Å on the RC because of the geometrical restraints from β-CD. The ligand is free from the geometrical restraints and can diffuse on the spherical space under the government of intermolecular interactions in windows above 7 Å and within 13 Å. Note that between 10 to 13 Å, the ligand can form favorable van der Waals (vdW) interactions if the ligand is far away from the X-axis and sticks to the outer surface of β-CD (Figure 7). Due to this reason, the ligand deviates from the artificial path along X-axis remarkably in that region (Figure 8). Apparently, even at the same COM distance of 10 Å, the ligand naturally tends to stay closer to β-CD when it is on the outer surface of β-CD with stronger attractions, than in the case where it is aligned to the X-axis, where no stronger attraction can be formed. This will certainly affect the shape of the PMF. When the RC is beyond 15 Å, the two molecules do not form strong interactions any more, and the ligand is totally free to diffuse in the spherical space in the simulation for one US window. With this concern, it is interesting to investigate how PMF will be affected by the *direct dissociation* along X-axis, and *indirect dissociation* where the ligand diffuses to the outer surface of β-CD and then dissociate.

**Figure 6.**
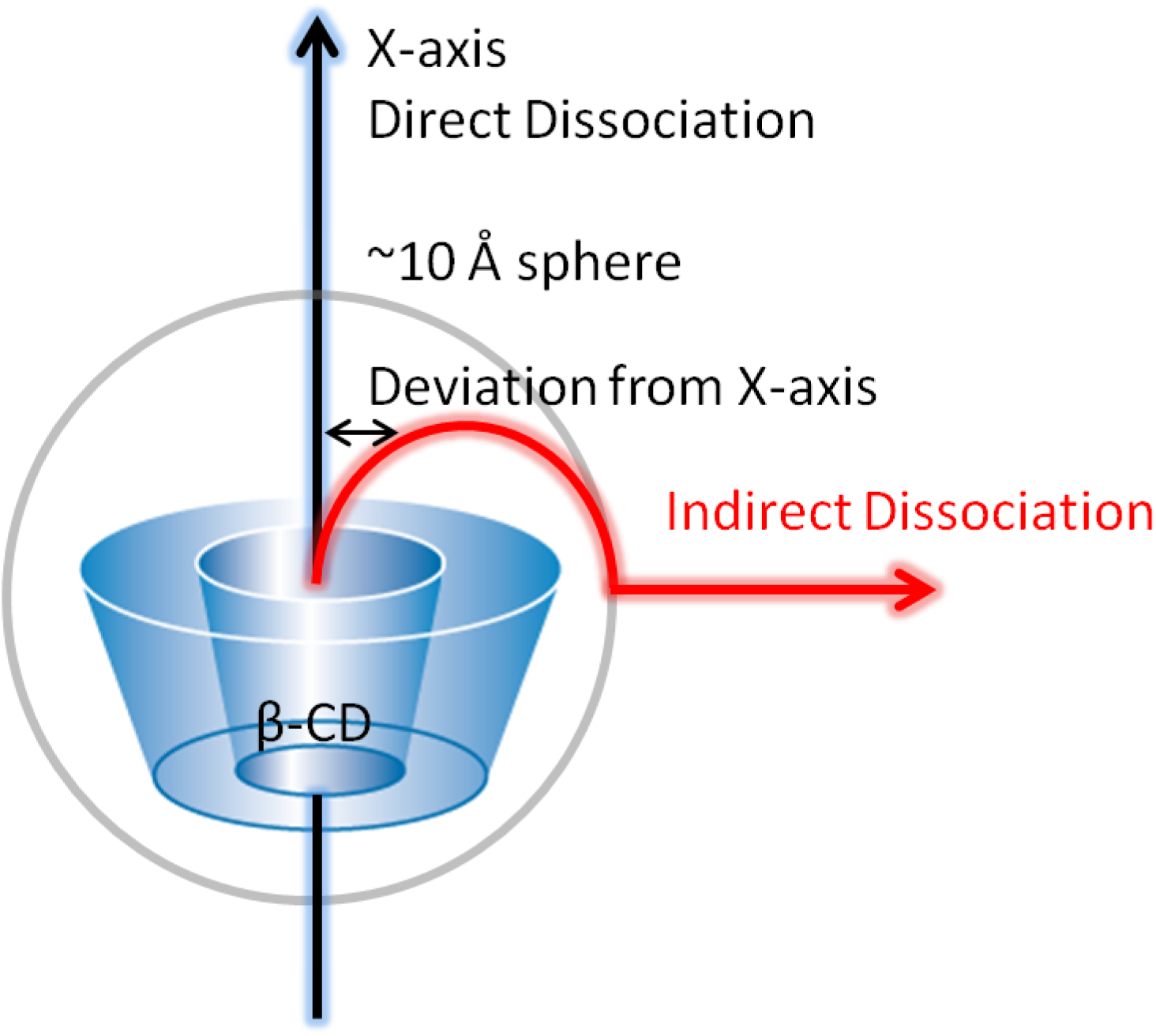
Direct and indirect ligand dissociation paths from β-CD. In direct dissociation, ligand moves out of binding site along X-axis. In indirect dissociation, ligand first diffuses to the outer surface of β-CD, which is about 10 Å from X-axis, then dissociate from there.

**Figure 7.**
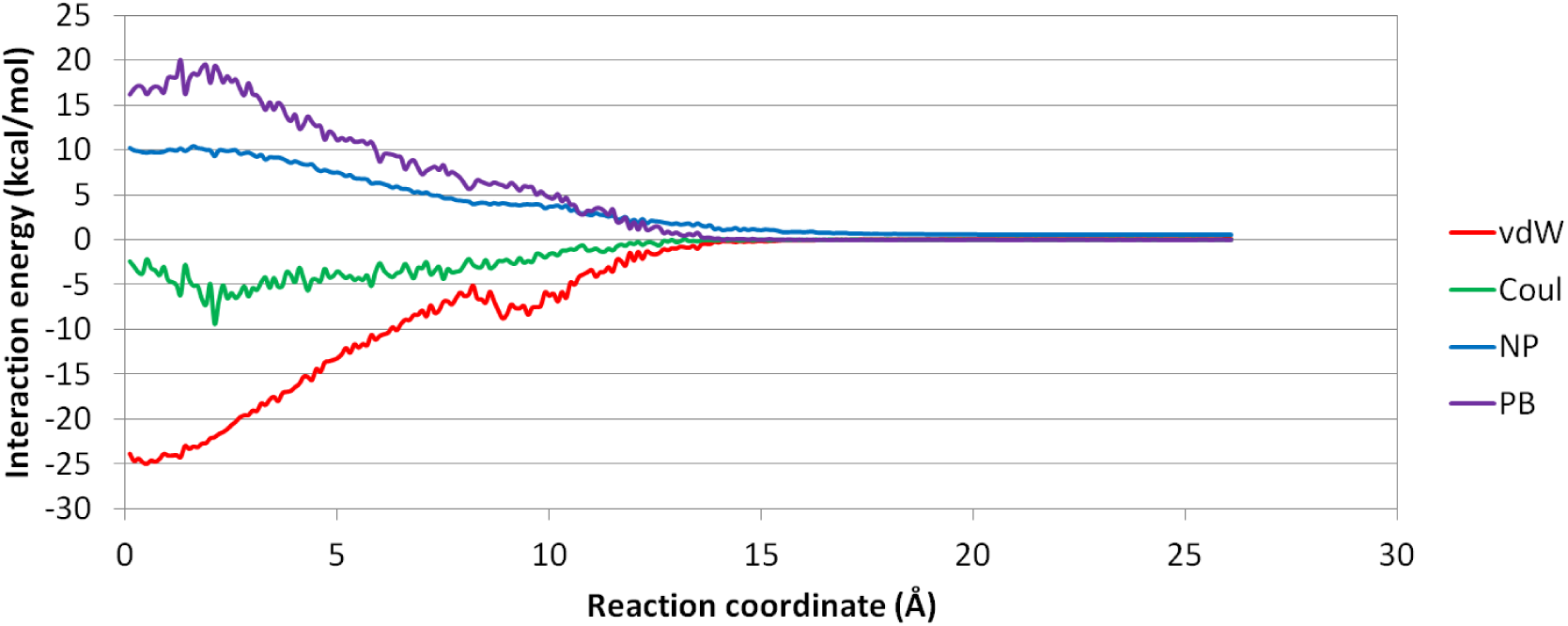
Plot of averaged MM/PBSA energy at each US window. Van der Waals energy (vdW), Coulombic energy (Coul), nonpolar solvation energy (NP) and PB solvation energy (PB) of aspirin from β-CD in Conf 1 along path A were averaged in the biased MD for each window.

**Figure 8.**
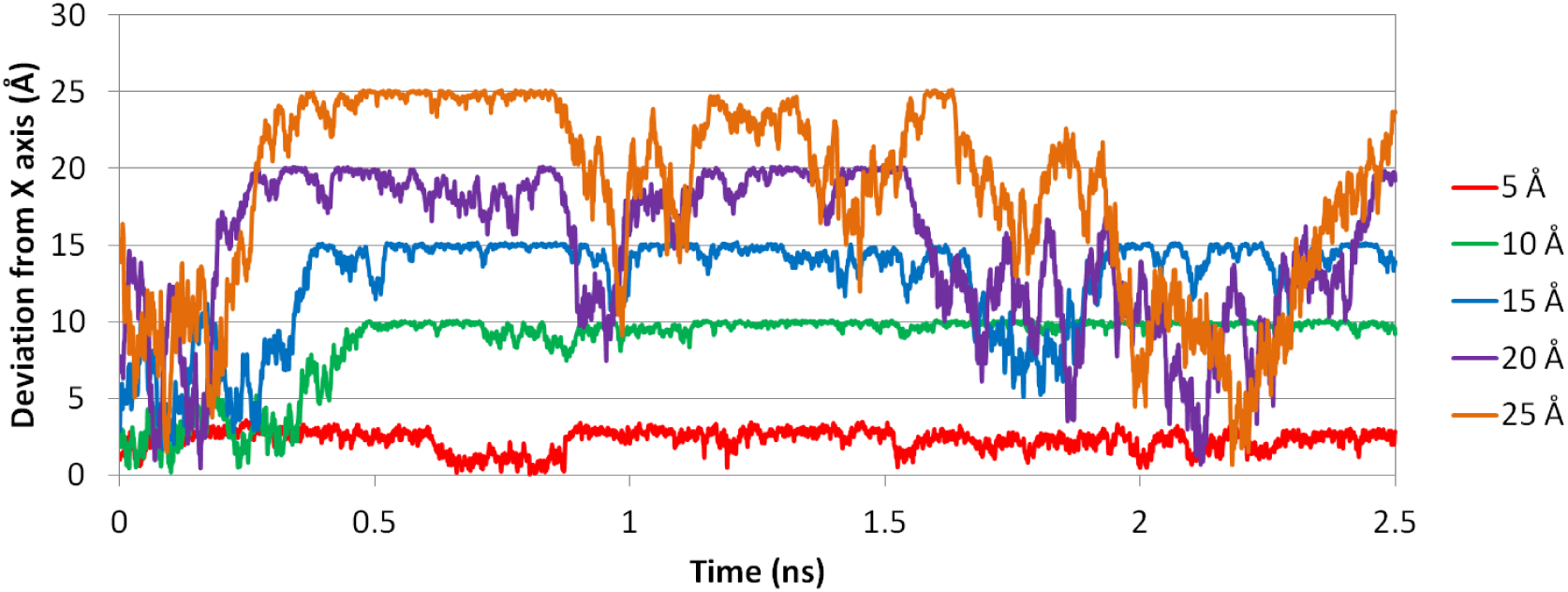
Deviation of aspirin from X axis at different RC distances. Deviations at 5, 10, 15, 20 and 25 Å during dissociation process of aspirin from β-CD in Conf 1 along path A are shown in red, green, cyan, purple and orange respectively.

According to data of aspirin deviation from X-axis (Figure 8), we generated a direct dissociation PMF and an indirect dissociation PMF by using the first 0.2 ns biased simulations and the last 2.0 ns biased simulations for aspirin respectively (Figure 9). As we anticipated, the direct PMF deviates from the indirect PMF that resembles the overall PMF. The direct PMF has a higher dissociation energy barrier at roughly 7 Å, and smooths out the energy valley at 10 Å. Compared to the artificially generated direct dissociation path along X-axis, the ligand is able to find the energy minimal path by walking on the surface of β-CD and depart from there in the indirect dissociation which reproduces the reality more, resulting in a lower PMF. The strong interaction between β-CD and aspirin through the outer surface of β-CD also generates the energy valley at 10 Å, which is missing in the direct PMF. To further explore how the initial guess of the dissociation path affects the PMF from US, we randomly picked data along RC from β-CD Conf 1, 2 and 3 to construct the cross PMF. In this way, we introduced conformational exchanges between biased MD simulation starting from Conf 1, 2 and 3. Not surprisingly, the cross PMF using data from Conf 1 and 3 is located between the original PMF curves (Figure 10). However, the dissociation energy barrier may deviate from 2.5 kcal/mol from data of Conf 3 to 4.4 kcal/mol from data of Conf 1. Considering the normal length of biased MD simulation in each window in US application is only on the scale of nanoseconds [11, 12], it is unlikely that such short biased simulation will thoroughly explore the conformational space restrained at the CV the users use to define the RC. Therefore, an initial guess of the dissociation path that deviates from the reality too much would not be brought back to the well-equilibrated state. This stands the red flag that when using US to compute PMF for a system, the initial guess of the dissociation path plays a crucial role, and non-energy-minimal dissociation path may lead to totally wrong PMF.

**Figure 9.**
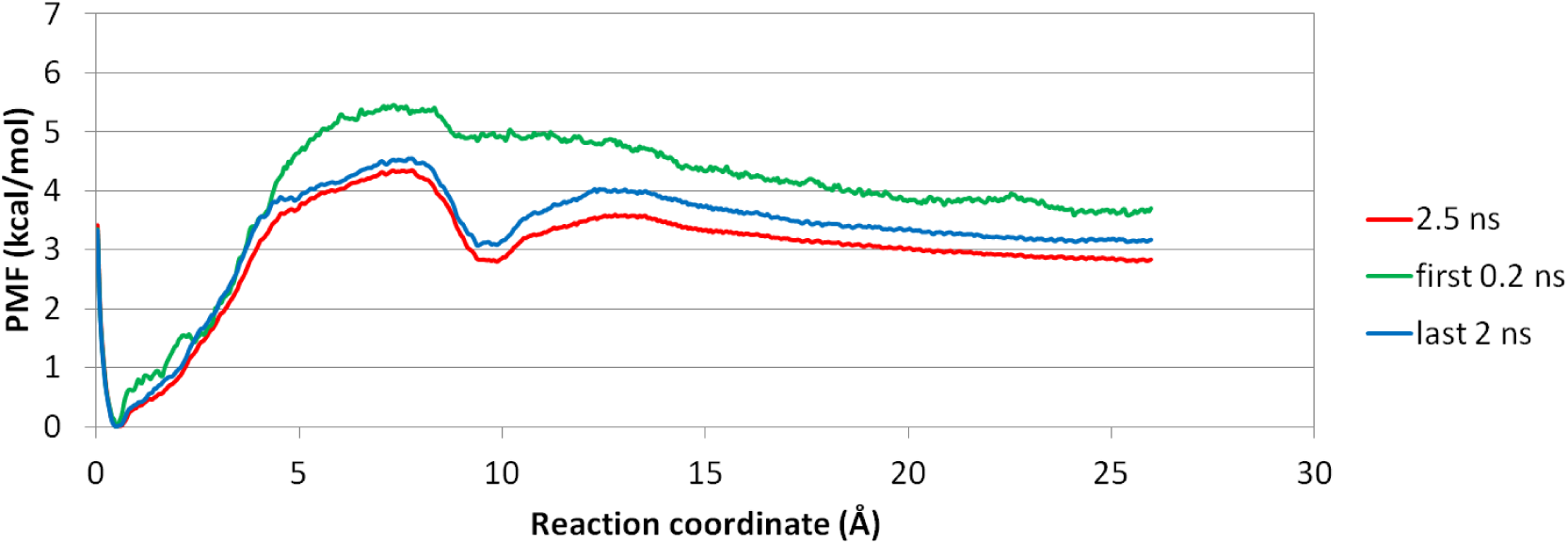
Three PMF plots of aspirin dissociation from β-CD in Conf 1 along path A. Red: using complete 2.5 ns biased MD for US. Green: using first 0.2 ns biased MD for US. Blue: using last 2.0 ns biased MD for US

**Figure 10.**
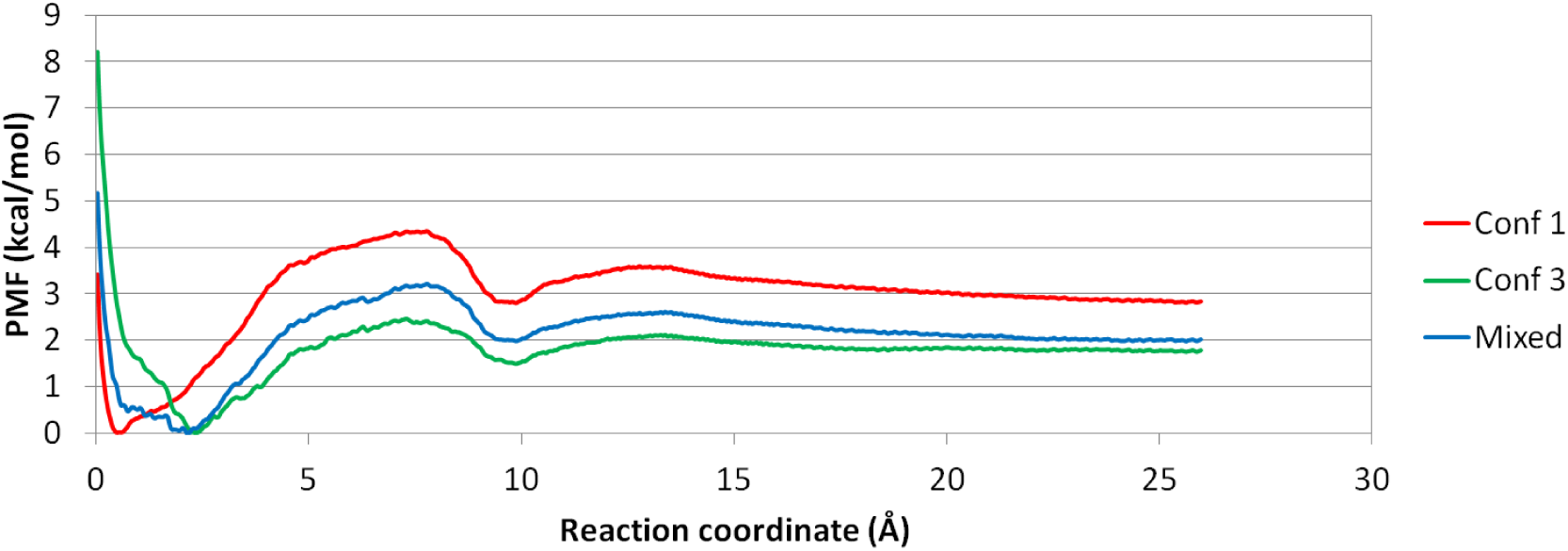
PMFs using Conf 1, Conf 3 and mixture of Conf 1 and 3 along path A. The mixed PMF in Conf 1 and 3 locates between the PMF of Conf 1 and 3.

### Unbinding process of p38α complex system

The unbinding process of a large protein-ligand system takes usually as long as microseconds or even days, involving slow protein motions, complicated ligand rearrangement and various non-bonding interactions. This is far beyond the practical timescale of MD simulations. Therefore, it is common to use enhanced sampling methods to sample the ligand dissociation pathways. Then US can be used to perform intensive sampling using such pathways as initial conformations to compute the PMF of dissociation. For p38α complex system, we prepared five paths obtained using AMD, SMD and manual pulling. Path 1 and 2 are dissociation of SB2 from DFG-in conformation using AMD equilibrated by conventional MD. One dissociation path of SB2 from DFG-out conformation was also generated in the same way. The directions of path 1 and 2 are slightly different, as indicated in Figure 11. Path 1 is a more straight forward dissociation where SB2 direct moves towards outside (Figure S4), path 2 is an indirect dissociation where SB2 adhered to the hinge region, and kept diffusing on surface of the hinge until eventually moving out. SB2 ligands in SMD and manual pulling are moving in between the two directions of path1 and 2.

PMF plots of the five paths of SB2 dissociation are shown in Figure 12. It is very interesting that except for the PMF from SMD, the other PMF plots predict similar binding affinities which are roughly 8 to 10 kcal/mol. The binding affinities from these paths somehow fall into the common range of drug-like compound binding affinities [59] and close to experimental value (Table 1), regardless of the diversity of behaviors of energy barriers along the dissociation pathway and protein conformations. US using SMD path, however, predicts a binding affinity of roughly 25 kcal/mol, which is incredibly large. The PMF from US is questionable. The nanosecond timescale biased MD simulation is incapable of exploring the entire sub conformational space of protein systems restraint at the specific CV or CVs used to represent the RC. Even for small systems like β-CD-ligand complexes, nanosecond level simulations fail to smooth out the effects from initial conformations of β-CD and it takes microsecond MD simulations to fully explore the conformational space of β-CD. Therefore, the binding affinity and energy barriers from the PMF computed using US are not undoubtedly reliable.

**Figure 11.**
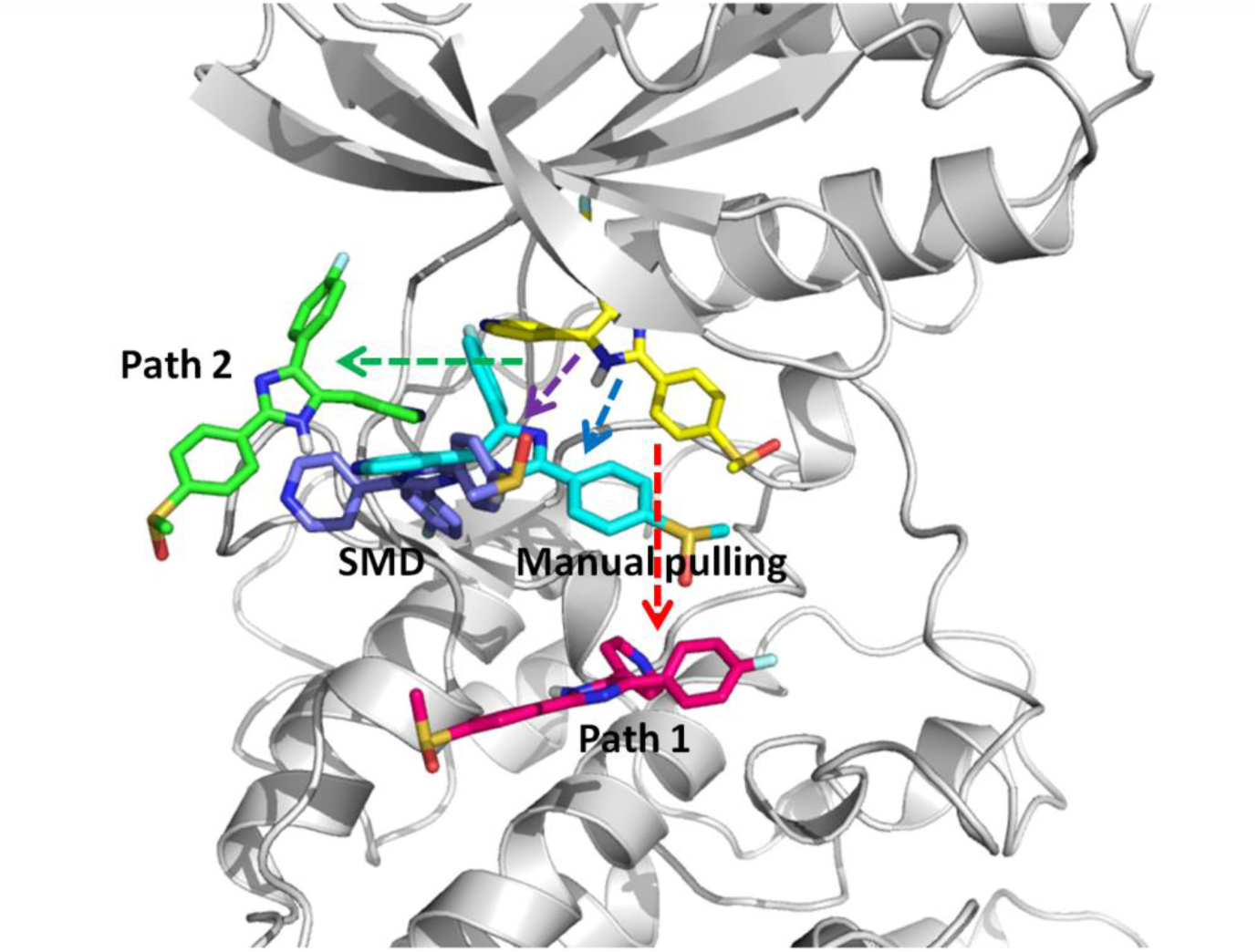
Four dissociation paths for SB2-p38α complex with DFG-in conformation. Yellow: crystal bound conformation of SB2. Red: path 1 where SB2 direct moves towards outside. Green: path 2 where SB2 diffuses on the surface of the hinge region until moving out. Cyan: manual pulling path. Purple: SMD path. SB2 in manual pulling and SMD paths moves out along directions in between the two directions of path1 and 2.

**Figure 12.**
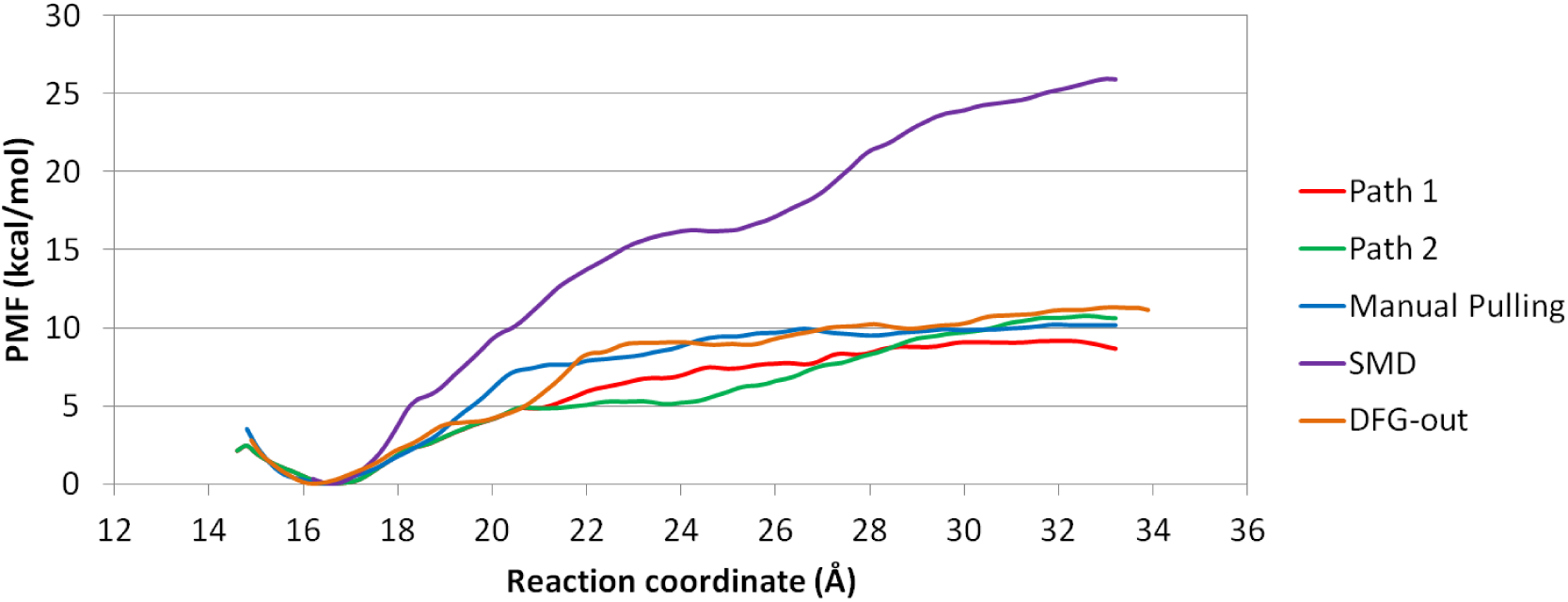
PMF plots of SB2-p38α complex. Red and green: path 1 and 2 from AMD path with conformation relaxed with short conventional MD. Cyan: manual pulling. Purple: SMD. Orange: AMD path with conformation relaxed with short conventional MD for dissociation of SB2 from DFG-out conformation.

US using paths from AMD predicts similar PMF. The binding affinities computed using path 1 and path 2 are 8.7 kcal/mol and 10.6 kcal/mol respectively, and are highly similar. In path 2, SB2 diffuses on surface of the hinge region during dissociation, which explains why its PMF reaches a low and flat region between 21 and 24 Å. In DFG-out conformation, after SB2 breaks its hydrogen bond with Met109, the 4-methylsulfinylphenyl group can rotate back inside cavity and its phenyl ring will form stacking interaction with Phe169 (Figure 13). However, the binding free energy of SB2 with DFG-out conformation from US is −11.2 kcal/mol, surprisingly similar to path 1 and 2 with DFG-in conformation, which agrees with previous NMR study [44] about the free conversion between DFG-in and DFG-out conformations of p38α while bound with SB2. This suggests that the PMF computed using conformations from dissociation paths yielded from AMD can consistently reproduce the binding affinity if the conformations along the dissociation paths are relaxed by conventional MD simulations, even if the MD simulations are short.

**Figure 13.**
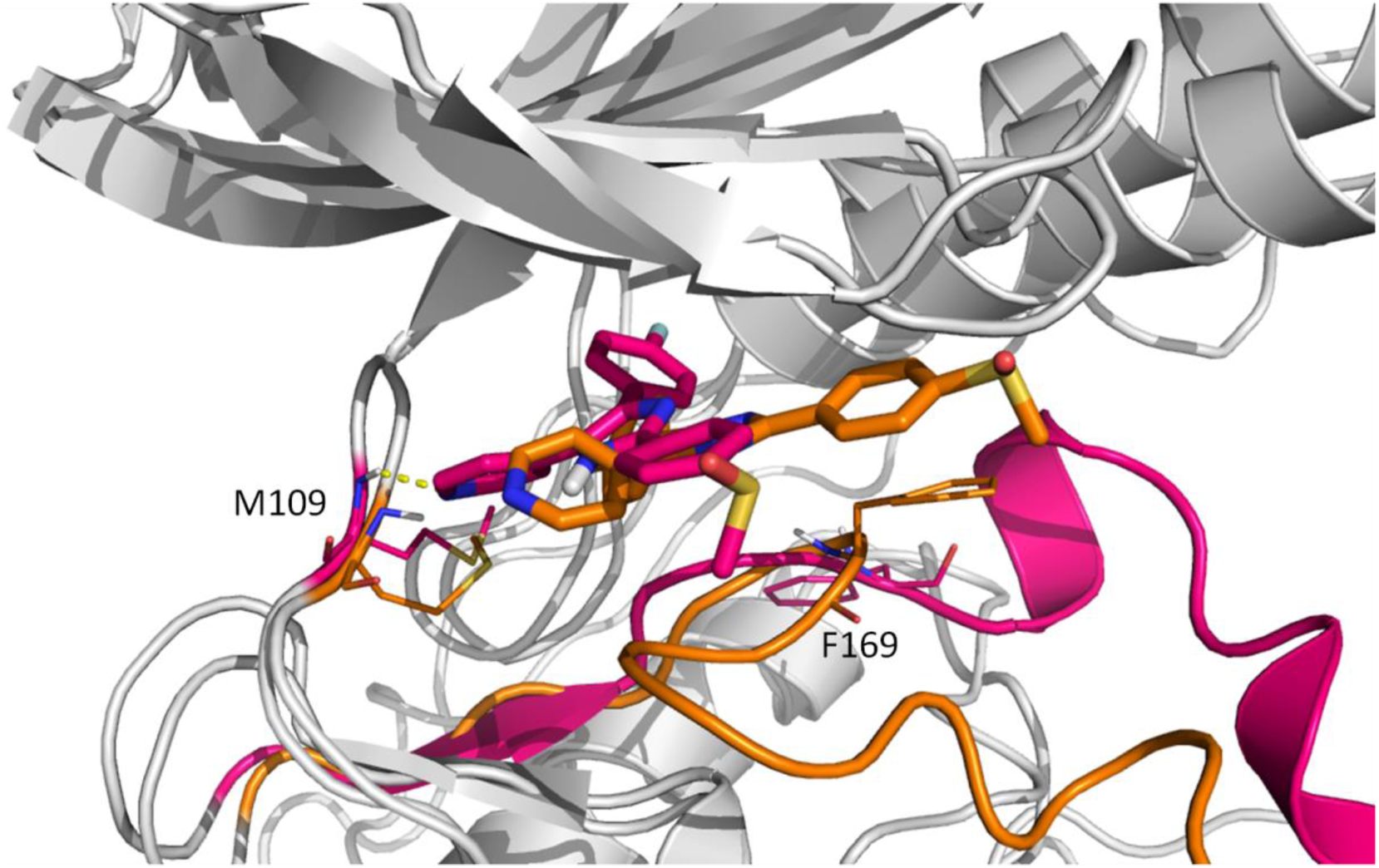
DFG-out path (orange color) VS path 1 (red color). During dissociation of SB2 from DFG-out conformation, 4-methylsulfinylphenyl group rotates back inside cavity and form stacking interaction with sidechain of Phe169. Hydrogen bonds between SB2 and Met109 are shown in dash line.

The different behaviors of protein motions and residue sidechain movement sampled from AMD, SMD and manual pulling may change the shape and height of PMF significantly. In free p38α crystal structure, Tyr35 has different orientation than SB2-p38α complex, instead of Tyr35 forming stacking interaction with the 4-methylsulfinylphenyl group of SB2, Tyr35 in free p38α rotates away and forms hydrogen bond with the sidechain of Arg67. Both free p38α crystal structure and our AMD simulation suggest that in order for SB2 to dissociate, Tyr35 will first rotates away into its position in free p38α crystal structure, breaking its stacking interaction with SB2, thus facilitating dissociation of SB2. However, in SMD, due to the enforced pulling force acted on SB2, Tyr35 doesn’t rotate away, but follows SB2 towards outside, after breaking stacking interaction with 4-methylsulfinylphenyl group at 24 Å, it immediately forms stacking interaction with fluorobenzene group of SB2, and doesn’t break until 29 Å (Figure 14). This cause much larger energy barrier comparing to other paths. Interestingly, manual pulling US doesn’t have this problem and actually performs quite similar to path 1 and 2, since all the residues of p38α are kept in the crystal structure positions, although Tyr35 doesn’t rotates away from SB2 at the beginning of dissociation, it doesn’t follow SB2 either, therefore avoiding unexpected interaction. Therefore, the initial guess of dissociation pathway for PMF calculation using US method must be validated and relaxed before performing biased sampling in each window for US.

**Figure 14.**
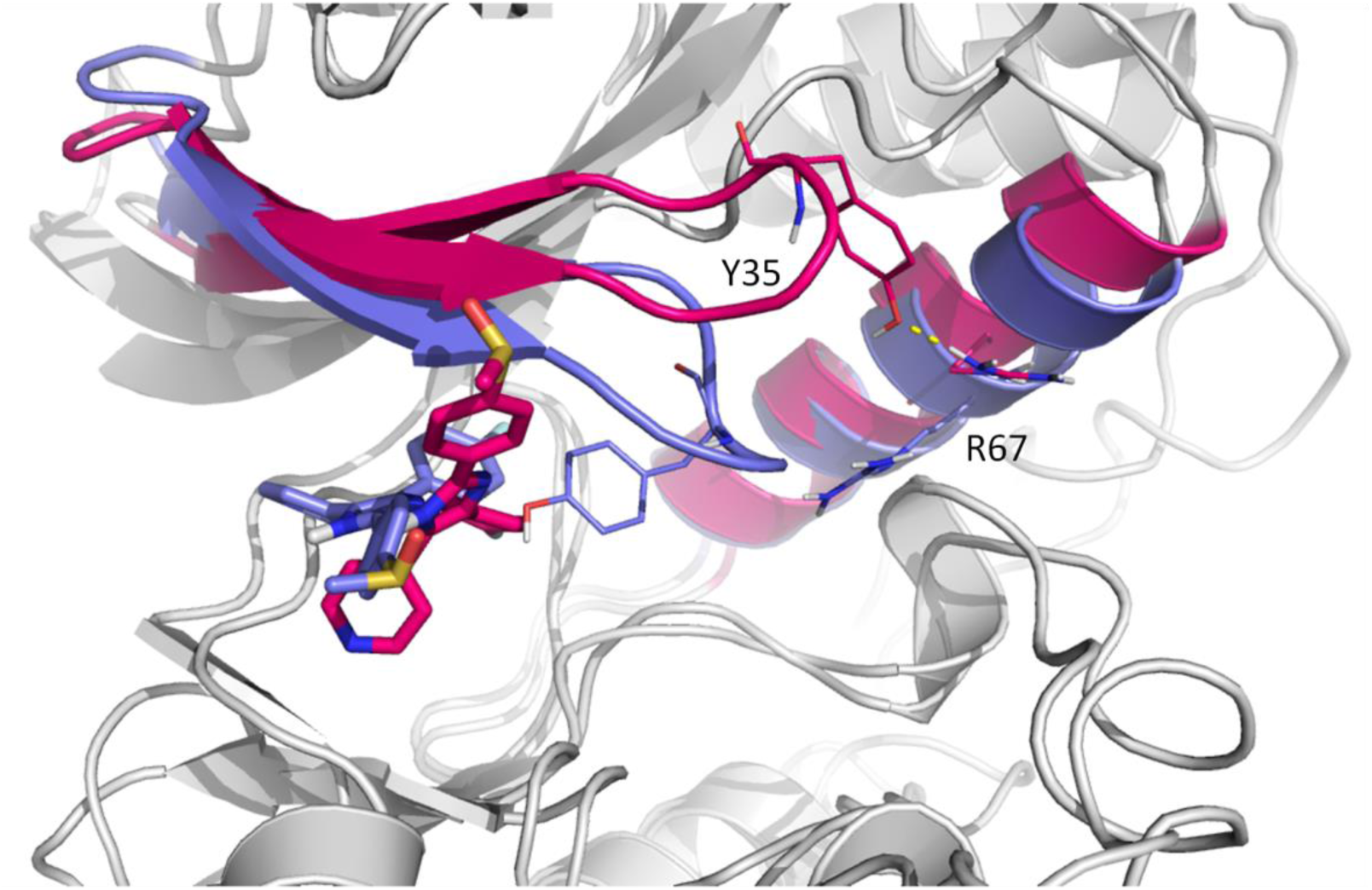
SMD path (purple color) VS path 1 (red color). Tyr35 rotates away in path 1 to form hydrogen bond (shown in dash line) with Arg67, and breaking stacking interaction with SB2, while Tyr35 in SMD follows SB2.

To confirm the lack of protein conformational change during dissociation process simulated by SMD and manual pulling simulations, we measured RMSD of backbone atoms of p38α in biased short simulations of US windows for path1, 2, manual pulling and SMD. Average RMSD were obtained in each US window with crystal structure of DFG-in p38α (1A9U) as reference (Figure S5). As expected, SMD and manual pulling simulations have smaller RMSD due to lack of necessary conformational change. SMD starts pulling after 100ns of conventional MD, which is enough to make the whole system reach equilibrium, but still not enough to get ready for dissociation. However, in researches using SMD to sample unbinding process of other kinase-ligand systems [60, 61], the lengths of conventional MD before SMD are mostly shorter than 100 ns, or even 10 ns, which might introduce artifacts into the simulation. Besides, previous research has shown that too large pulling speed or pulling force constant in SMD would result in instability of system or significantly higher energy barrier [62]. Therefore, future dissociation sampling with SMD should be more rigorous with system setting up and conformation selection to minimize artificial errors. Gladly, several methods have been brought up to improve free energy estimates and conformation sampling of non-equilibrium SMD simulation. Based on the forward–reverse method, Nategholeslam et al. introduced bin-passing method to better separate of the reversible and irreversible work distributions, and achieve faster convergence [63]. By combining configurational freezing and nonuniform particle-selection scheme, Riccardo Chelli developed local sampling in steered Monte Carlo simulations that can enhance the accuracy of the free energy calculation [64]. Whalen et al. applied hybrid steered MD-docking method to better rank inhibitor affinities against a flexible drug target [65].

## Conclusion

We applied US to investigate the dissociation processes of β-CD and p38α complex systems with several conformational sampling methods, including AMD, SMD and manual pulling. We also investigated the influences of the computed PMF from conformations, dissociation pathways, intensity of the biased sampling in US and dissociation pathway sampling method. Different β-CD conformations can change the depth of PMF by more than 50%, and this suggest that nanosecond timescale biased simulation is unable to remove the effects from initial conformations, even for small host like β-CD. By comparing the direct and indirect pathway of β-CD complex dissociation, we also found that different dissociation paths can result in appearance and disappearance of local minima, and non-energy-minimal dissociation path may lead to wrong PMF. Commonly used enhanced sampling methods that provide initial conformations for US were discussed. For large protein-ligand system, SMD can efficiently pull ligand out of binding cavity, however, the artificial force may introduce unexpected interactions or miss necessary conformational change in sampling of dissociation process, thus requiring careful setting up and screening of structures. Manual pulling is similar to SMD, but may be suitable for sampling dissociation path of system that not requiring much host conformational change. Compared to these two methods, universal acceleration sampling method like AMD is able to simulate adequate details along the dissociation pathway. By relaxing the conformations along dissociation pathways sampled by AMD using short MD simulations, we obtained more reliable initial conformations for US, leading to consistent and reliable PMF. Therefore, we suggest that US can be a very reliable method for computing PMF, if a proper enhanced sampling method is used and the initial conformations are properly relaxed to remove the bias from the enhanced technique.

## Abbreviations

US: ,umbrella sampling
MD: molecular dynamics
AMD: accelerated molecular dynamics
SMD: steered molecular dynamics
β-CD: β-cyclodextrin
PMF: potential of mean force
RC: reaction coordinate
CV: collective variable
COM: center of mass
RMSD: root mean square deviation

---

Supporting Materials for

## Artifact of Restraints

Due to the limitation of distance restraint algorithm used in AMBER, when COM of ligand was close to 0 Å on X axis, where COM of β-CD was located, ligand could jump back and forth between path A and B. The harmonic potential added to restrain the distance between β-CD and ligand was irrelevant from directions and therefore causing an issue shown in Figure S3. When the distance between COMs of β-CD and ligand was larger than 1 Å, the problem disappeared because the gap had become too large for ligand to jump through. We ran multiple runs for RC smaller than 1 Å to get a rough closer PMF. The future version of restraint setting in AMBER may consider restraining vector instead distance, so this issue can be revisited and solved. It’s also noted that for the 0 Å on X axis, where the COMs of β-CD and ligand were overlapped, the peak split into two located on both sides. It’s an artifact of restraint setting in AMBER, which caused the unusual high energy barrier at 0 Å position in Figure 5.

**Figure S1.**
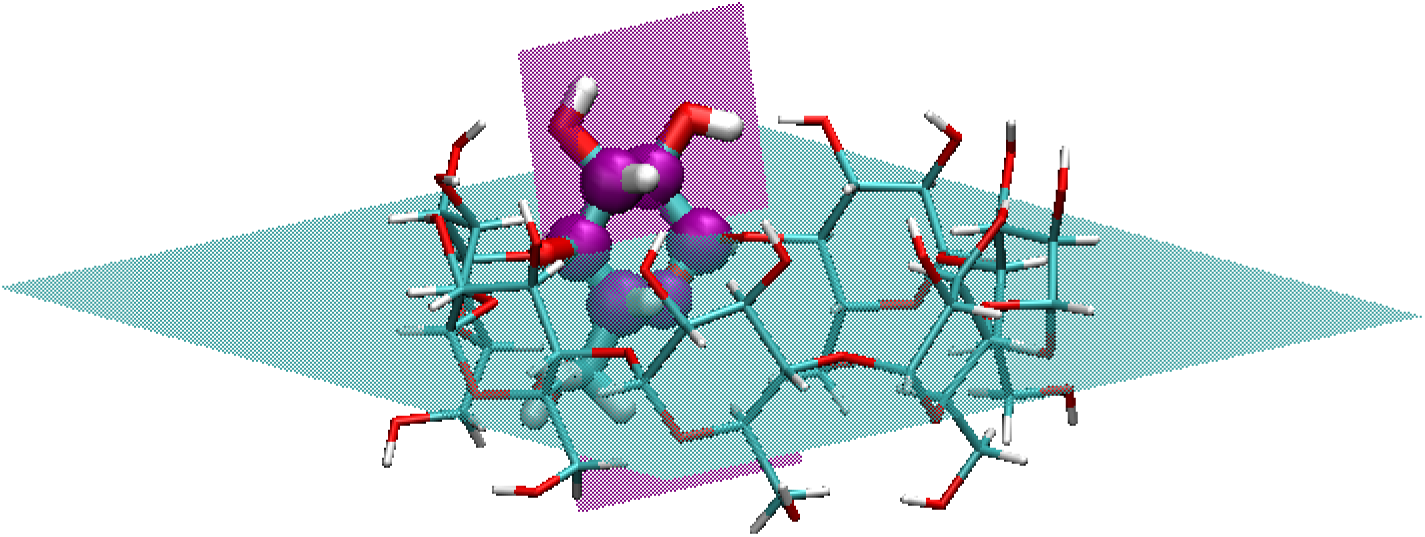
Representation of one fingerprint angle in β-CD. The regression plane of entire molecule is in shown in cyan color and the regression plane of the six atoms highlighted by purple balls in one glucose unit is shown in purple color. The fingerprint angle of one glucose unit is defined by the dihedral angle between the two regression planes.

**Figure S2.**
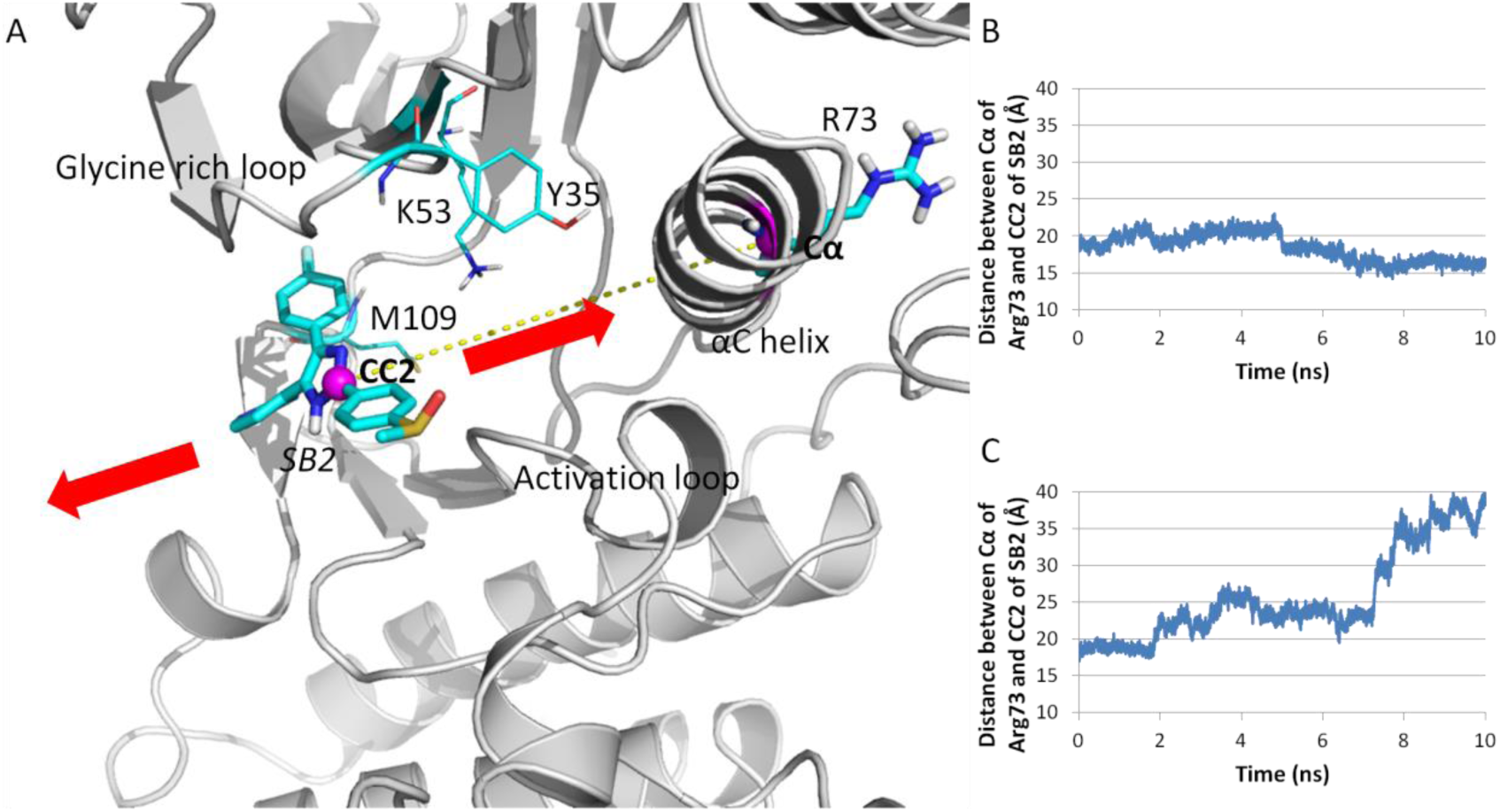
Reconstruction of dissociation path from AMD. Path 1 is built from AMD path that conformational relaxed by two 10 ns conventional MD. (A) SB2 in one of the two 10 ns conventional MD moves towards inside the cavity, while SB2 in the other conventional MD moves towards outside. Arg73 and SB2 are shown in bold licorice structure, Cα of Arg73 and CC2 of SB2 are indicated by purple ball structure, other interacting residues are shown in thin licorice structure. (B) SB2 moving inside the cavity indicated by the decreasing distance between SB2 and Arg73. (C) SB2 moving outside the cavity indicated by the increasing distance between SB2 and Arg73.

**Figure S3.**
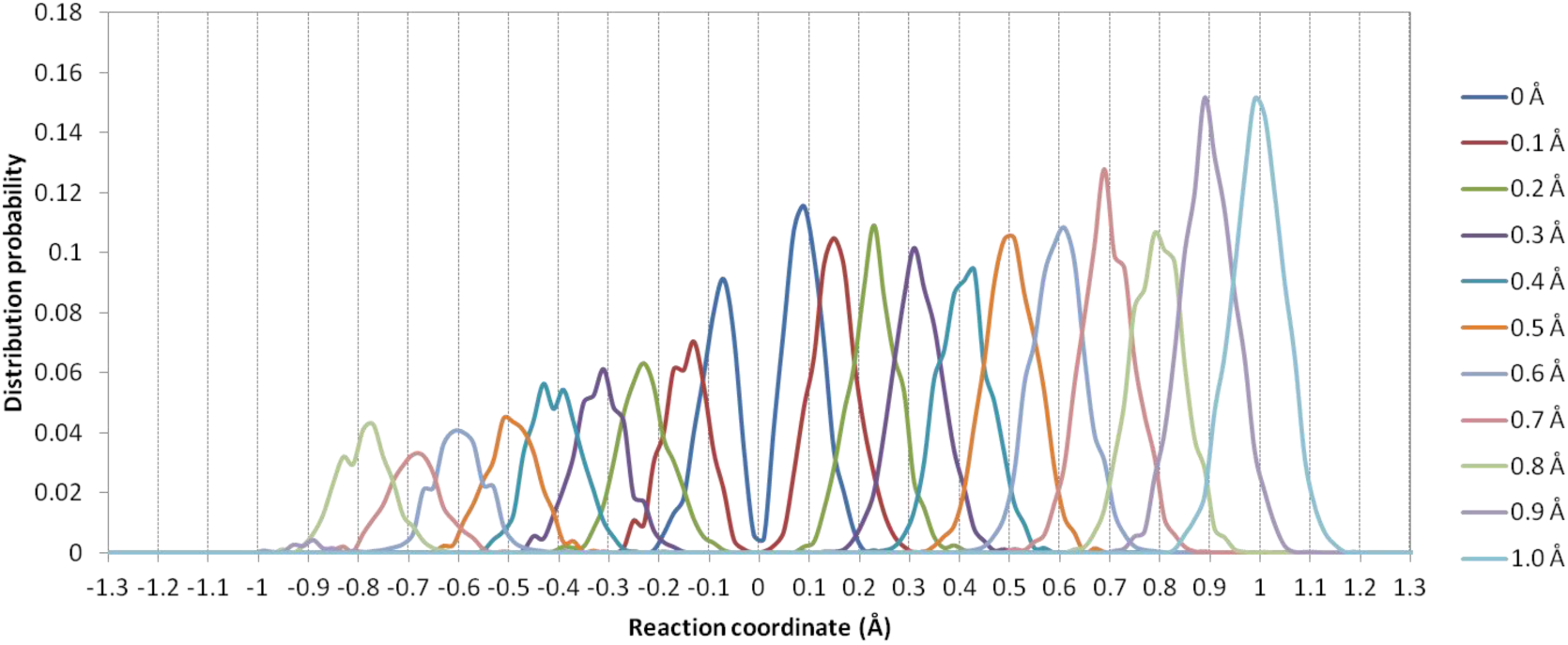
Selected distribution probability of ligand in US along path A. When distance of COMs of aspirin and β-CD is smaller than 1 Å, ligand jumps back and forth between path A and B during dissociation process of aspirin from β-CD in Conf 1 along path A, ligand stops jumping when RC distance reaches 1 Å.

**Figure S4.**
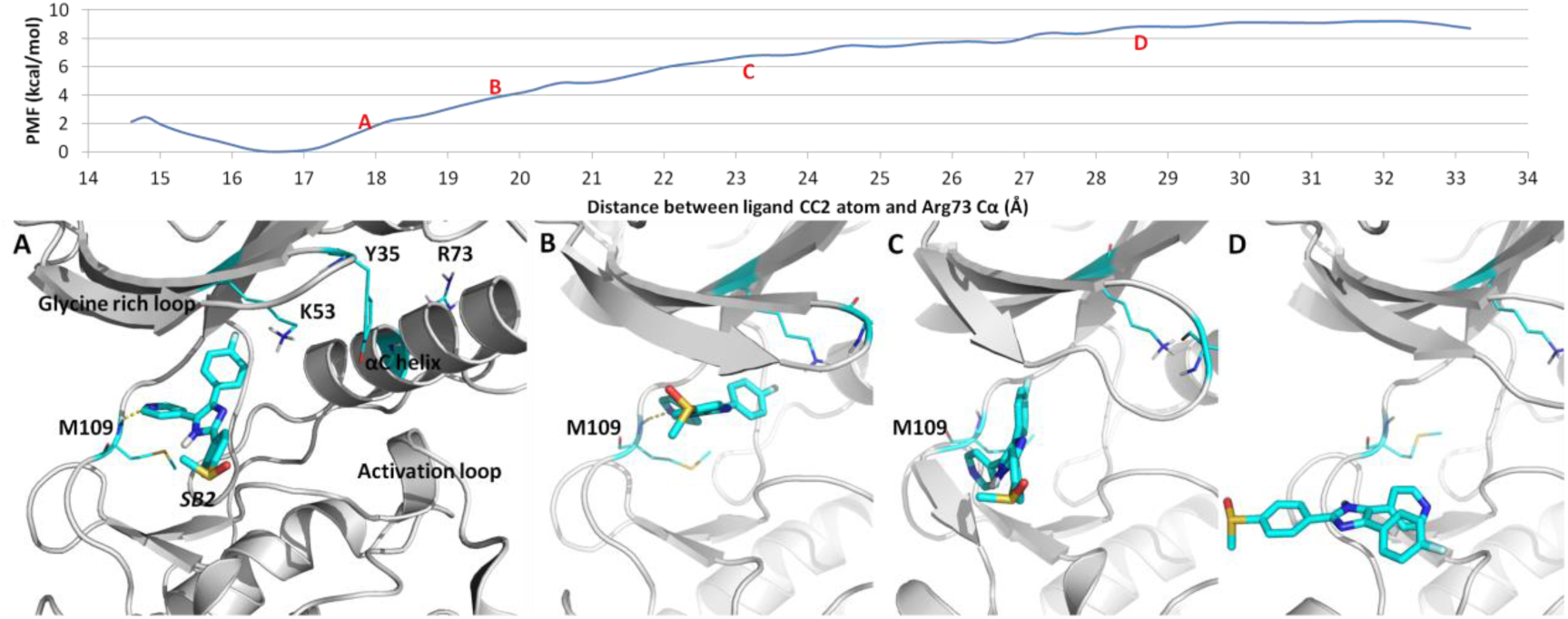
PMF of path 1 from US and the selected snapshots during dissociation. Hydrogen bonds between SB2 and p38α are shown in dash line. (A) SB2 breaks hydrogen bond with Lys53 side-chain and stacking interaction with Tyr35. (B) 4-methylsulfinylphenyl group of SB2 starts diffusing towards outside the cavity (C) fluorophenyl ring of SB2 moves out of the hydrophobic pocket and hydrogen bond between pyridine nitrogen and Met109 breaks. (D) SB2 is outside the edge of binding cavity.

**Figure S5.**
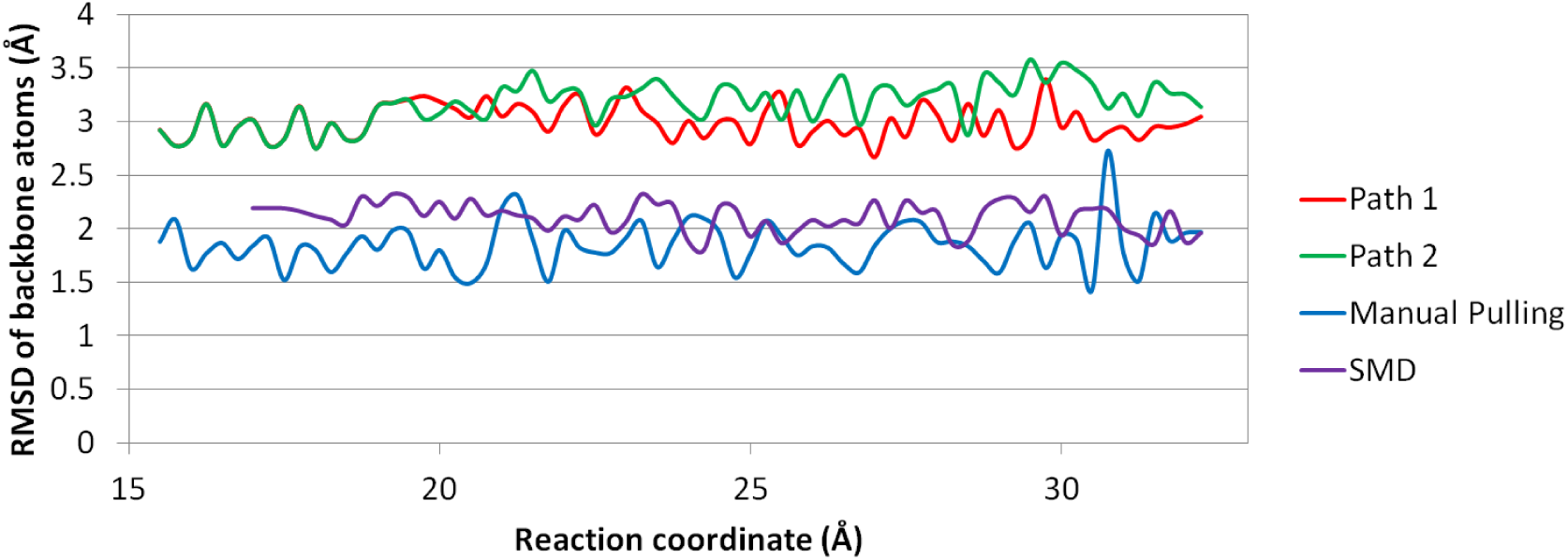
Plot of averaged RMSD of backbone atoms of p38α at each US window. RMSD values of backbone atoms of p38α from path1, 2, manual pulling and SMD paths are averaged in the biased MD for each window. Path 1 and 2 show higher RMSD due to AMD sampling more protein conformational change than SMD and manual pulling.

## References

1. Beveridge DL, DiCapua FM. Free energy via molecular simulation: applications to chemical and biomolecular systems. Annual review of biophysics and biophysical chemistry. 1989;18:431–92. Epub 1989/01/01. doi:10.1146/annurev.bb.18.060189.002243. PubMed PMID: 2660832.

2. Foloppe N, Hubbard R. Towards Predictive Ligand Design With Free-Energy Based Computational Methods?. Current medicinal chemistry. 2006;13(29):3583–608. doi: http://dx.doi.org/10.2174/092986706779026165.

3. Kollman P. Free energy calculations: Applications to chemical and biochemical phenomena. Chemical Reviews. 1993;93(7):2395–417. doi:10.1021/cr00023a004.

4. Zwanzig RW. HIGH-TEMPERATURE EQUATION OF STATE BY A PERTURBATION METHOD .1.NONPOLAR GASES. Journal of Chemical Physics. 1954;22(8):1420–6. PubMed PMID: WOS:A1954UC16200028.

5. Khavrutskii IV, Wallqvist A. Improved Binding Free Energy Predictions from Single-Reference Thermodynamic Integration Augmented with Hamiltonian Replica Exchange. Journal of chemical theory and computation. 2011;7(9):3001–11. doi:10.1021/ct2003786.

6. Torrie GM, Valleau JP. NON-PHYSICAL SAMPLING DISTRIBUTIONS IN MONTE-CARLO FREE-ENERGY ESTIMATION - UMBRELLA SAMPLING. Journal of Computational Physics. 1977;23(2):187–99. doi:10.1016/0021-9991(77)90121-8. PubMed PMID: WOS:A1977CX19800007.

7. Hainam D, Hirst JD, Wheatley RJ. Calculation of Partition Functions and Free Energies of a Binary Mixture Using the Energy Partitioning Method: Application to Carbon Dioxide and Methane. Journal of Physical Chemistry B. 2012;116(15):4535–42. doi:10.1021/jp212168f. PubMed PMID: WOS:000302924800016.

8. D L Beveridgea, DiCapua FM. Free Energy Via Molecular Simulation: Applications to Chemical and Biomolecular Systems. Annual review of biophysics and biophysical chemistry. 1989;18(1):431–92. doi:10.1146/annurev.bb.18.060189.002243. PubMed PMID: 2660832.

9. Chipot C. Frontiers in free-energy calculations of biological systems. Wiley Interdisciplinary Reviews: Computational Molecular Science. 2014;4(1):71–89. doi:10.1002/wcms.1157.

10. Vashisth H, Skiniotis G, Brooks CL. Collective Variable Approaches for Single Molecule Flexible Fitting and Enhanced Sampling. Chemical Reviews. 2014;114(6):3353–65. doi:10.1021/cr4005988.

11. Velez-Vega C, Gilson MK. Force and Stress along Simulated Dissociation Pathways of Cucurbituril-Guest Systems. Journal of chemical theory and computation. 2012;8(3):966–76. Epub 2012/07/04. doi:10.1021/ct2006902. PubMed PMID: 22754402; PubMed Central PMCID: PMCPmc3383817.

12. Sun H, Tian S, Zhou S, Li Y, Li D, Xu L, et al. Revealing the favorable dissociation pathway of type II kinase inhibitors via enhanced sampling simulations and two-end-state calculations. Scientific reports. 2015;5:8457. Epub 2015/02/14. doi:10.1038/srep08457. PubMed PMID: 25678308; PubMed Central PMCID: PMCPmc4326958.

13. Li W, Shen J, Liu G, Tang Y, Hoshino T. Exploring coumarin egress channels in human cytochrome P450 2A6 by random acceleration and steered molecular dynamics simulations. Proteins. 2011;79(1):271–81. Epub 2010/11/09. doi:10.1002/prot.22880. PubMed PMID: 21058395.

14. Marzinek JK, Bond PJ, Lian G, Zhao Y, Han L, Noro MG, et al. Free Energy Predictions of Ligand Binding to an α-Helix Using Steered Molecular Dynamics and Umbrella Sampling Simulations. Journal of Chemical Information and Modeling. 2014;54(7):2093–104. doi:10.1021/ci500164q.

15. Nguyen H, Le L. Steered molecular dynamics approach for promising drugs for influenza A virus targeting M2 channel proteins. European biophysics journal: EBJ. 2015;44(6):447–55. Epub 2015/06/03. doi:10.1007/s00249-015-1047-4. PubMed PMID: 26033540.

16. Rodriguez RA, Yu L, Chen LY. Computing Protein-Protein Association Affinity with Hybrid Steered Molecular Dynamics. Journal of chemical theory and computation. 2015;11(9):4427–38. Epub 2015/09/15. doi:10.1021/acs.jctc.5b00340. PubMed PMID: 26366131; PubMed Central PMCID: PMCPmc4565455.

17. Sun H, Li Y, Tian S, Wang J, Hou T. P-loop conformation governed crizotinib resistance in G2032R-mutated ROS1 tyrosine kinase: clues from free energy landscape. PLoS computational biology. 2014;10(7):e1003729. Epub 2014/07/18. doi:10.1371/journal.pcbi.1003729. PubMed PMID: 25033171; PubMed Central PMCID: PMCPmc4102447.

18. Sun H, Chen P, Li D, Li Y, Hou T. Directly Binding Rather than Induced-Fit Dominated Binding Affinity Difference in (S)- and (R)-Crizotinib Bound MTH1. Journal of chemical theory and computation. 2016;12(2):851–60. Epub 2016/01/15. doi:10.1021/acs.jctc.5b00973. PubMed PMID: 26764587.

19. Doudou S, Burton NA, Henchman RH. Standard Free Energy of Binding from a One-Dimensional Potential of Mean Force. Journal of chemical theory and computation. 2009;5(4):909–18. Epub 2009/04/14. doi:10.1021/ct8002354. PubMed PMID: 26609600.

20. Gumbart JC, Roux B, Chipot C. Standard binding free energies from computer simulations: What is the best strategy?. Journal of chemical theory and computation. 2013;9(1):794–802. Epub 2013/06/25. doi:10.1021/ct3008099. PubMed PMID: 23794960; PubMed Central PMCID: PMCPmc3685508.

21. Woo HJ, Roux B. Calculation of absolute protein-ligand binding free energy from computer simulations. Proceedings of the National Academy of Sciences of the United States of America. 2005;102(19):6825–30. Epub 2005/05/04. doi:10.1073/pnas.0409005102. PubMed PMID: 15867154; PubMed Central PMCID: PMCPmc1100764.

22. Jo S, Suh D, He Z, Chipot C, Roux B. Leveraging the Information from Markov State Models To Improve the Convergence of Umbrella Sampling Simulations. The journal of physical chemistry B. 2016;120(33):8733–42. Epub 2016/07/14. doi:10.1021/acs.jpcb.6b05125. PubMed PMID: 27409349.

23. Breslow R, Dong SD. Biomimetic Reactions Catalyzed by Cyclodextrins and Their Derivatives. Chem Rev. 1998;98(5):1997–2012. Epub 2002/02/19. PubMed PMID: 11848956.

24. Challa R, Ahuja A, Ali J, Khar RK. Cyclodextrins in drug delivery: an updated review. AAPS PharmSciTech. 2005;6(2):E329–57. Epub 2005/12/16. doi:10.1208/pt060243. PubMed PMID: 16353992; PubMed Central PMCID: PMCPmc2750546.

25. Davis ME, Brewster ME. Cyclodextrin-based pharmaceutics: past, present and future. Nat Rev Drug Discov. 2004;3(12):1023–35. Epub 2004/12/02. doi:10.1038/nrd1576. PubMed PMID: 15573101.

26. Del Valle EMM. Cyclodextrins and their uses: a review. Process Biochemistry. 2004;39(9):1033–46. doi: http://dx.doi.org/10.1016/S0032-9592(03)00258-9.

27. Lai WF. Cyclodextrins in non-viral gene delivery. Biomaterials. 2014;35(1):401–11. Epub 2013/10/10. doi:10.1016/j.biomaterials.2013.09.061. PubMed PMID: 24103652.

28. Marchetti L, Levine M. Biomimetic Catalysis. ACS Catalysis. 2011;1(9):1090–118. doi:10.1021/cs200171u.

29. Singh M, Sharma R, Banerjee UC. Biotechnological applications of cyclodextrins. Biotechnology advances. 2002;20(5-6):341–59. Epub 2003/10/11. PubMed PMID: 14550020.

30. Barros TC, Stefaniak K, Holzwarth JF, Bohne C. Complexation of Naphthylethanols with β-Cyclodextrin. The Journal of Physical Chemistry A. 1998;102(28):5639–51. doi:10.1021/jp9803844.

31. Fukahori T, Kondo M, Nishikawa S. Dynamic Study of Interaction between β-Cyclodextrin and Aspirin by the Ultrasonic Relaxation Method. The Journal of Physical Chemistry B. 2006;110(9):4487–91. doi:10.1021/jp058205n.

32. Izatt RM, Pawlak K, Bradshaw JS, Bruening RL. Thermodynamic and Kinetic Data for Macrocycle Interaction with Cations, Anions, and Neutral Molecules. Chemical Reviews. 1995;95(7):2529–86. doi:10.1021/cr00039a010.

33. Nilsson M, Valente AJM, Olofsson G, Söderman O, Bonini M. Thermodynamic and Kinetic Characterization of Host-Guest Association between Bolaform Surfactants and α- and β-Cyclodextrins. The Journal of Physical Chemistry B. 2008;112(36):11310–6. doi:10.1021/jp802963x.

34. Nishikawa S, Ugawa T, Fukahori T. Molecular Recognition Kinetics of β-Cyclodextrin for Several Alcohols by Ultrasonic Relaxation Method. The Journal of Physical Chemistry B. 2001;105(31):7594–7. doi:10.1021/jp010535u.

35. Rekharsky MV, Inoue Y. Complexation Thermodynamics of Cyclodextrins. Chem Rev. 1998;98(5):1875–918. Epub 2002/02/19. PubMed PMID: 11848952.

36. Schneider, H-J, Hacket F, Rüdiger V, Ikeda H. NMR Studies of Cyclodextrins and Cyclodextrin Complexes. Chemical Reviews. 1998;98(5):1755–86. doi:10.1021/cr970019t.

37. Yim CT, Zhu XX, Brown GR. Kinetics of Inclusion Reactions of β-Cyclodextrin with Several Dihydroxycholate Ions Studied by NMR Spectroscopy. The Journal of Physical Chemistry B. 1999;103(3):597–602. doi:10.1021/jp9833909.

38. Adams JL, Badger AM, Kumar S, Lee JC. p38 MAP kinase: molecular target for the inhibition of pro-inflammatory cytokines. Progress in medicinal chemistry. 2001;38:1–60. Epub 2002/01/05. PubMed PMID: 11774793.

39. Cuenda A, Rousseau S. p38 MAP-kinases pathway regulation, function and role in human diseases. Biochimica et biophysica acta. 2007;1773(8):1358–75. Epub 2007/05/08. doi:10.1016/j.bbamcr.2007.03.010. PubMed PMID: 17481747.

40. Kumar S, Boehm J, Lee JC. p38 MAP kinases: key signalling molecules as therapeutic targets for inflammatory diseases. Nature reviews Drug discovery. 2003;2(9):717–26. Epub 2003/09/03. doi:10.1038/nrd1177. PubMed PMID: 12951578.

41. Peifer C, Wagner G, Laufer S. New approaches to the treatment of inflammatory disorders small molecule inhibitors of p38 MAP kinase. Current topics in medicinal chemistry. 2006;6(2):113–49. Epub 2006/02/04. PubMed PMID: 16454763.

42. Schindler JF, Monahan JB, Smith WG. p38 pathway kinases as anti-inflammatory drug targets. Journal of dental research. 2007;86(9):800–11. Epub 2007/08/28. PubMed PMID: 17720847.

43. Huse M, Kuriyan J. The conformational plasticity of protein kinases. Cell. 2002;109(3):275–82. Epub 2002/05/23. PubMed PMID: 12015977.

44. Vogtherr M, Saxena K, Hoelder S, Grimme S, Betz M, Schieborr U, et al. NMR characterization of kinase p38 dynamics in free and ligand-bound forms. Angewandte Chemie (International ed in English). 2006;45(6):993–7. Epub 2005/12/24. doi:10.1002/anie.200502770. PubMed PMID: 16374788.

45. Cezard C, Trivelli X, Aubry F, Djedaini-Pilard F, Dupradeau, F-Y. Molecular dynamics studies of native and substituted cyclodextrins in different media: 1. Charge derivation and force field performances. Physical Chemistry Chemical Physics. 2011;13(33):15103–21. doi:10.1039/C1CP20854C.

46. Pedretti A, Villa L, Vistoli G. VEGA--an open platform to develop chemo-bio-informatics applications, using plug-in architecture and script programming. Journal of computer-aided molecular design. 2004;18(3):167–73. Epub 2004/09/17. PubMed PMID: 15368917.

47. Frisch MJ, Trucks GW, Schlegel HB, Scuseria GE, Robb MA, Cheeseman JR, et al. Gaussian 09. Wallingford, CT, USA: Gaussian, Inc.; 2009.

48. Wang Z, Canagarajah BJ, Boehm JC, Kassisa S, Cobb MH, Young PR, et al. Structural basis of inhibitor selectivity in MAP kinases. Structure (London, England: 1993). 1998;6(9):1117–28. Epub 1998/10/01. PubMed PMID: 9753691.

49. Simard JR, Getlik M, Grutter C, Pawar V, Wulfert S, Rabiller M, et al. Development of a fluorescent-tagged kinase assay system for the detection and characterization of allosteric kinase inhibitors. Journal of the American Chemical Society. 2009;131(37):13286–96. Epub 2009/07/04. doi:10.1021/ja902010p. PubMed PMID: 19572644.

50. Casper D, Bukhtiyarova M, Springman EB. A Biacore biosensor method for detailed kinetic binding analysis of small molecule inhibitors of p38alpha mitogen-activated protein kinase. Analytical biochemistry. 2004;325(1):126–36. Epub 2004/01/13. PubMed PMID: 14715293.

51. Hamelberg D, Mongan J, McCammon JA. Accelerated molecular dynamics: a promising and efficient simulation method for biomolecules. J Chem Phys. 2004;120(24):11919–29. Epub 2004/07/23. doi:10.1063/1.1755656. PubMed PMID: 15268227.

52. Jensen MO, Park S, Tajkhorshid E, Schulten K. Energetics of glycerol conduction through aquaglyceroporin GlpF. Proceedings of the National Academy of Sciences of the United States of America. 2002;99(10):6731–6. Epub 2002/05/09. doi:10.1073/pnas.102649299. PubMed PMID: 11997475; PubMed Central PMCID: PMCPmc124471.

53. Torrie GM, Valleau JP. Monte Carlo free energy estimates using non-Boltzmann sampling: Application to the sub-critical Lennard-Jones fluid. Chemical Physics Letters. 1974;28(4):578–81. doi: http://dx.doi.org/10.1016/0009-2614(74)80109-0.

54. Torrie GM, Valleau JP. Nonphysical sampling distributions in Monte Carlo free-energy estimation: Umbrella sampling. Journal of Computational Physics. 1977;23(2):187–99. doi: http://dx.doi.org/10.1016/0021-9991(77)90121-8.

55. Case DA, Babin V, Berryman JT, Betz RM, Cai Q, Cerutti DS, et al. {Amber 14}2014.

56. Souaille M, Roux Bt. Extension to the weighted histogram analysis method: combining umbrella sampling with free energy calculations. Computer physics communications. 2001;135(1):40–57. doi: http://dx.doi.org/10.1016/S0010-4655(00)00215-0.

57. Still WC, Tempczyk A, Hawley RC, Hendrickson T. SEMIANALYTICAL TREATMENT OF SOLVATION FOR MOLECULAR MECHANICS AND DYNAMICS. Journal of the American Chemical Society. 1990;112(16):6127–9. doi:10.1021/ja00172a038. PubMed PMID: WOS:A1990DR56800038.

58. Jorgensen WL, Chandrasekhar J, Madura JD, Impey RW, Klein ML. Comparison of simple potential functions for simulating liquid water. The Journal of Chemical Physics. 1983;79(2):926–35. doi:10.1063/1.445869.

59. Keseru GM, Makara, GM. The influence of lead discovery strategies on the properties of drug candidates. Nat Rev Drug Discov.2009;8(3):203–12. doi: http://www.nature.com/nrd/journal/v8/n3/suppinfo/nrd2796_S1.html.

60. Capelli AM, Costantino G. Unbinding pathways of VEGFR2 inhibitors revealed by steered molecular dynamics. J Chem Inf Model. 2014;54(11):3124–36. Epub 2014/10/10. doi:10.1021/ci500527j. PubMed PMID: 25299731.

61. Yang L-J, Zou J, Xie H-Z, Li L-L, Wei Y-Q, Yang, S-Y. Steered Molecular Dynamics Simulations Reveal the Likelier Dissociation Pathway of Imatinib from Its Targeting Kinases c-Kit and Abl. PLOS ONE. 2009;4(12):e8470. doi:10.1371/journal.pone.0008470.

62. Zhang Z, Santos AP, Zhou Q, Liang L, Wang Q, Wu T, et al. Steered molecular dynamics study of inhibitor binding in the internal binding site in dehaloperoxidase-hemoglobin. Biophysical Chemistry. 2016;211:28–38. doi: http://dx.doi.org/10.1016/j.bpc.2016.01.003.

63. Nategholeslam M, Gray CG, Tomberli B. Implementation of the forward-reverse method for calculating the potential of mean force using a dynamic restraining protocol. The journal of physical chemistry B. 2014;118(49):14203–14. Epub 2014/11/06. doi:10.1021/jp504942t. PubMed PMID: 25372312.

64. Chelli R. Local Sampling in Steered Monte Carlo Simulations Decreases Dissipation and Enhances Free Energy Estimates via Nonequilibrium Work Theorems. Journal of chemical theory and computation. 2012;8(11):4040–52. Epub 2012/11/13. doi:10.1021/ct300348w. PubMed PMID: 26605571.

65. Whalen KL, Chang KM, Spies MA. Hybrid Steered Molecular Dynamics-Docking: An Efficient Solution to the Problem of Ranking Inhibitor Affinities Against a Flexible Drug Target. Molecular informatics. 2011;30(5):459–71. Epub 2011/07/09. doi:10.1002/minf.201100014. PubMed PMID: 21738559; PubMed Central PMCID: PMCPmc3129543.

